# Unconventional mRNA processing and degradation pathways for the polycistronic y*rzI* mRNA in *B. subtilis*

**DOI:** 10.1101/2024.11.26.625382

**Authors:** Laetitia Gilet, Magali Leroy, Alexandre Maes, Ciarán Condon, Frédérique Braun

**Author notes:** These authors contributed equally (Laetitia Gilet, Magali Leroy).

## Abstract

The <u>r</u>ibosome-<u>a</u>ssociated <u>e</u>ndoribonuclease Rae1 cleaves the *Bacillus subtilis yrzI* operon mRNA in a translation-dependent manner. This operon encodes up to four small peptides, S1027, YrzI, S1025 and S1024, whose functions were unknown. We show that YrzI is toxic at high concentrations, but co-expression with S1025 abolishes its toxicity. We show that a highly stable mRNA species containing the YrzI and S1025 open reading frames results from endoribonucleolytic cleavage upstream of the *yrzI* ORF followed by 5’-exoribonucleolytic processing arrested by ribosomes bound to its exceptionally strong Shine-Dalgarno sequence. Degradation of this mRNA requires either translation-dependent cleavage within S1025 by Rae1 or direct attack from the structured 3’-end by 3’-exoribonucleases. Both pathways are atypical for a *B. subtilis* mRNA.

## Introduction

Two major mRNA degradation pathways have been characterized in *Bacillus subtilis* thus far [1,2]. The 5’-exoribonucleolytic (5’-exo) pathway resembles the mRNA decapping and Xrn1-dependent pathway found in eukaryotes [1,2] After deprotection of the mRNA by an RNA pyrophosphohydrolase, *e.g.* BsRppH, the 5’-exoribonuclease complex RNase J1/J2 degrades the mRNA all the way to the 3’ end [3]. In the endoribonucleolytic (endo) pathway, the mRNA is cleaved internally and the resulting fragments degraded by either 3’- or 5’-exoribonucleases. The key endoribonucleases known to initiate mRNA decay in *B. subtilis* are the single strand-specific enzyme RNase Y, a functional analog of *E. coli* RNase E, and the double strand-specific enzyme RNase III [4]. In general, the activity of these enzymes is inhibited by translation, with the flow of ribosomes occluding access to cleavage sites within open reading frames (ORFs) [5–9]. In 2017, we identified a novel component of the *B. subtilis* mRNA decay machinery, the ribosome associated endoribonuclease Rae1 [10] that, in contrast to RNase III and RNase Y, requires translation for cleavage [11,12].

We have thus far identified two Rae1 targets: the *bmrBCD* operon mRNA [12], encoding a multidrug transporter, and the *yrzI* operon mRNA [10]. Ribosome profiling experiments have suggested that the latter operon, in addition to encoding the annotated 49-amino acid (aa) YrzI peptide, expresses three supplementary peptides S1027, S1025 and S1024, that are 38, 17 and 52 aa’s in length, respectively [13]. We previously showed that Rae1 cleaves within the S1025 ORF in a translation-dependent manner, confirming that the 17-aa S1025 peptide is indeed translated [10]. However, the functions of the different peptides expressed from this operon remain mysterious.

Here, we show that the overexpression of the YrzI peptide impairs *B. subtilis* growth, an effect that is counteracted by S1025, suggesting that these two peptides constitute a toxin-antitoxin (TA) system. We also show that expression of this operon is governed at the transcriptional level by the transition state regulator, AbrB, and at the post-transcriptional level via non-canonical mRNA processing and degradation pathways. Two primary transcripts undergo processing to a short highly stable ~500 nt mRNA fragment that is protected from 5’-degradation by ribosomes bound to an exceptionally strong Shine-Dalgarno (SD) sequence located upstream of *yrzI*. With the 5’-degradation pathway blocked by initiating ribosomes, elimination of this mRNA fragment requires degradation by one of two alternative pathways (i) translation-dependent endoribonucleolytic cleavage by Rae1, followed by degradation of the unprotected upstream and downstream fragments by 3’- and 5’-exoribonucleases or (ii) direct attack by 3’-exoribonucleases from the 3’ end of the mRNA despite the presence of a protective transcription terminator.

## Materials and Methods

### Strain construction

Oligonucleotides and strains used in the paper are listed in Tables S1 and S2, respectively.

The *rph*::spc, *rnr*::tc, *yhaM*::pm, *pnp*::kan and *pnp::cm* cassettes to make strains CCB329, CCB395, CCB396, CCB407, CCB409 and CCB1210 in our lab background (SSB1002 = W168 *trpC*+) were a kind gift from David Bechhofer, and have been described previously [31]. Strain CC376 was constructed by transferring the *rae1*::pMUTIN construct from strain CCB375 into CCB396. Strain CC761 was constructed by transferring the *rny::spec* construct from strain CCB441 into CCB375.

The plasmid over-expressing the YrzI peptide was constructed by amplifying the *yrzI* gene by PCR using oligos CC1731/1732 and cloned in pDG148 cleaved with SalI and HindIII. The resulting plasmid pDGYrzI (pl. 687) contains the *yrzI* gene under *Pspac* control, followed by an *E. coli* rRNA transcription terminator to limit transcription of downstream plasmid DNA. This plasmid was transformed into CCB375 (*Δrae1)* to create strain CCB815. We initially failed to transform this plasmid into wild-type cells. We thus re-isolated it from strain CCB815, transformed it with high efficiency into SSB1002 (WT) to create strain CCB839. The plasmid was re-isolated from CCB839 to confirm its sequence. pDGYrzI-S1025 (pl. 726) and pDGS1025 (pl. 727) were constructed by amplifying *yrzI-*S1025 and S1025 fragments by PCR using oligo pairs CC1731/1809 and CC1808/1809. PCR fragments were cleaved with SalI and HindIII and cloned in pDG148. The resulting plasmids were transferred to *B. subtilis* WT and *Δrae1* cells to create strains CCB913, CCB914 (pl. 726) and CCB915, CCB916 (pl. 727), respectively.

### Northern blots and primer extension assays

Northern blots were performed on total RNAs isolated either by the glass beads/phenol method described in [32] or by the RNAsnap method described in [33]. Northern blots were performed as described previously [4]. Primer extension assays were performed using the oligo CC1589 on glass bead/phenol extracted RNAs as described previously [34].

### Spot dilution assays

Cells were grown in 2xYT medium to OD_600_ = 0.6. When cells contained either the empty plasmid pDGYrzI (pl687), pDGYrzI-S1025 (pl726) and pDGS1025 (pl727), serial dilutions were spot on LB plates containing kanamycin (5 µg/ml) to maintain the plasmid. When indicated, IPTG at 100µM to induce the expression of the inserted gene was added.

### Ribosomal Subunit (30S) Protection Assay

Five nanomoles of a 3’-labelled (p^32^Cp) *yrzI* transcript was mixed with 500 nM *B. subtilis* 30S ribosomal subunits isolated from an RNase J1-depleted strain as described in [34] in 4 μl RNase J1 reaction buffer. The *yrzI* transcript was synthesized *in vitro* by T7 RNA polymerase using a MegaShortScript kit (Ambion) from PCR fragments amplified from chromosomal DNA using oligo pair CC3176/CC3042 (CC3176 had an integrated T7 RNA polymerase promoter sequence). Subunits were allowed to bind for 10 min at 37°C, followed by 10 min on ice. A total of 1.8 μg (1 μl) RNase J1 was added to start the reaction. Reactions were incubated at 25°C for 10 min, stopped as above, and loaded directly onto 5% polyacrylamide denaturing gels.

## Results

### YrzI encodes a toxic peptide that is counteracted by S1025

The *yrzI* operon encodes two proteins (YrhF and YrhG) and four small peptides S1027, YrzI, S1025 and S1024, whose functions are all unknown (Fig. 1A). We previously showed that Rae1 cleaves within the *S1025* ORF in a translational-dependent manner between codons 13 and 14 to initiate the degradation of the *yrzI* polycistronic mRNA and that deletion of the *rae1* gene led to an accumulation of three major transcripts of ~2.4 kb (P1-T4), ~0.8 kb (P3-T4) and ~0.5 kb in size (R-T4), all containing the *yrzI* open reading frame [10] (Fig. 1A). Since small peptides can have toxic effects on bacterial cell growth [14], we wondered whether the Rae1 cleavage site within S1025 might serve to eliminate this mRNA to protect cells from potential toxicity of one of its encoded peptides. Because YrzI was the originally annotated peptide of this operon and we had direct evidence that S1025 was translated, we began by analyzing the effect of these two peptides on cell growth by constructing two replicative plasmids, pDG-YrzI and pDG-S1025, permitting over-expression of YrzI and S1025 under control of an IPTG-dependent promoter. Induction of *yrzI* expression by addition of IPTG led to a 3 to 4-log inhibitory effect on *B. subtilis* growth in WT and *Δrae1* cells (Fig. 1B), suggesting that the YrzI peptide is indeed toxic at high doses. In contrast, no toxic effect of S1025 over-expression was observed in either WT or *Δrae1* cells (Fig. 1B). Since many toxins are encoded in operons with their antitoxins just downstream [14], we assayed the effect of producing both peptides together by cloning the *YrzI* and *S1025* region under control of an IPTG-dependent promoter (pDG-YrzI-S1025). Remarkably, co-expression of S1025 abolished the toxicity of YrzI (Fig. 1B). The protective effect of adding S1025 is not simply due to destabilisation of the *yrzI*-*S1025* mRNA through addition of the Rae1 cleavage site, since expression of S1025 neutralized the toxicity of YrzI even in absence of Rae1 (Fig. 1B). Thus, two different levels of controlling YrzI expression/toxicity exist: cleavage of its mRNA by Rae1 within the *S1025* ORF and a Rae1-independent antitoxin effect of the S1025 peptide.

**Figure 1.**
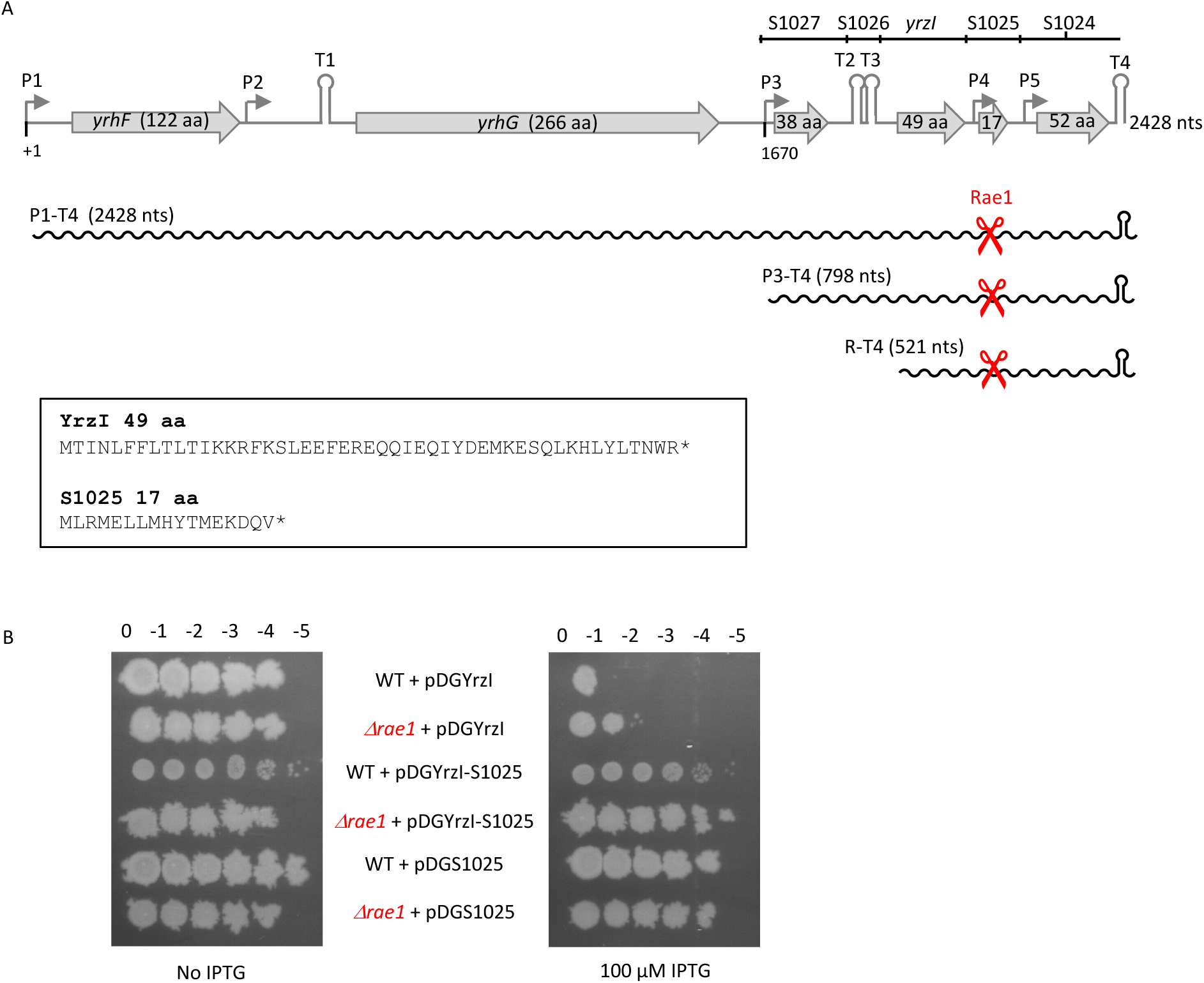
The toxicity of YrzI is counteracted by S1025. (A) Structure of the *yrzI* operon. Putative ORFs are represented by grey arrows, the putative promoters by rightward-pointing arrows and putative transcription terminators by the hairpin structures and Rae1 cleavage site within *S1025* by scissors. The size of the ORFs is given in amino acids (aa). The transcripts from this locus are shown as wavy lines. The sequence of the YrzI and S1025 peptide is shown. (B) Ten-fold serial dilution of WT and *Δrae1* strains containing the YrzI, S1025 or YrzI+S1025 overexpression plasmids pDGYrzI, pDGS1025 and pDGYrzI-S1025, respectively, spotted on LB plates with or without IPTG (100 µM).

### The polycistronic *yrzI* mRNA is regulated by AbrB at the transcriptional level

It has previously been shown that the expression of the *bmrBCD operon* is repressed by the transition state regulator AbrB at the transcriptional level [15], in addition to the post-transcriptional regulation mediated by Rae1[12]. Interestingly, transcriptional repression by AbrB is also predicted to be shared by the *yrzI* operon [16] and an AbrB binding site was mapped upstream of the P3 promoter [17]. To confirm this, we measured the levels of *yrzI* mRNA in strains lacking *rae1*, *abrB* or both, at mid-exponential growth phase, by Northern blot (Fig. 2). Deletion of *abrB* alone caused a small accumulation of the P3-T4 *yrzI* transcript compared to WT (Fig. 2, compare lane 3 to 1), and deletion of both *rae1* and *abrB* caused greater increase in accumulation of both P3-T4 and R-T4 transcripts than deletion of *rae1* alone (Fig. 2, compare lane 4 to 2). These results confirm that AbrB is a repressor of *yrzI* operon expression in mid-exponential phase. Expression of the *yrzI* operon is thus modulated at two levels, transcriptionally by AbrB and post-transcriptionally by Rae1.

**Figure 2.**
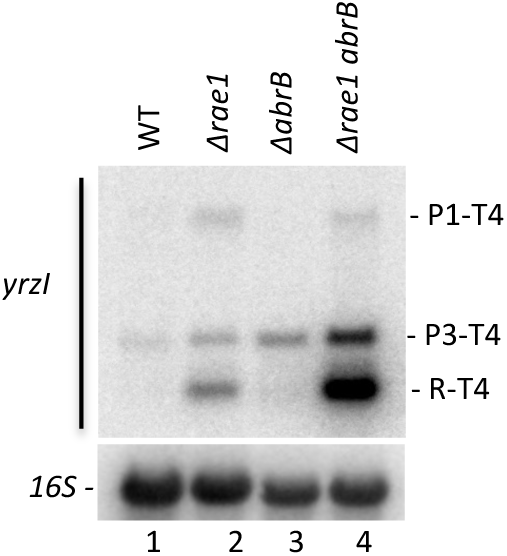
The polycistronic *yrzI* mRNA is regulated by AbrB at the transcriptional level. Northern blot showing steady state level of the *yrzI* and *abrB* transcripts in WT, *Δrae1*, *ΔabrB* and *Δrae1 ΔabrB* strains at mid-log phase (OD600 = 0.5). The blot was probed with oligo CC1589 (*yrzI*) and with a probe complementary to 16S rRNA (CC058) as a loading control.

### Mapping of the major transcripts around the *yrzI* locus

A transcriptome study by [18], identified several potential promoter (P1, P2, P3, P4 and P5) and terminator sequences (T2, T3 and T4) around the *yrzI* locus (Fig. 1A). However, Northern blots probed for *yrzI* only accounted for transcripts originating at P1 or P3 and ending at T4 (Fig. 3A, left panel). To get a more complete picture of the transcripts mapping to this locus, we hybridized Northern blots with oligonucleotide probes complementary to the *S1027* or *S1024* ORFs, upstream and downstream of *yrzI*, respectively. To determine their sensitivity to Rae1, these blots contained total RNA isolated from WT, *Δrae1* and plasmid complemented strains grown in the absence or presence of IPTG.

**Figure 3.**
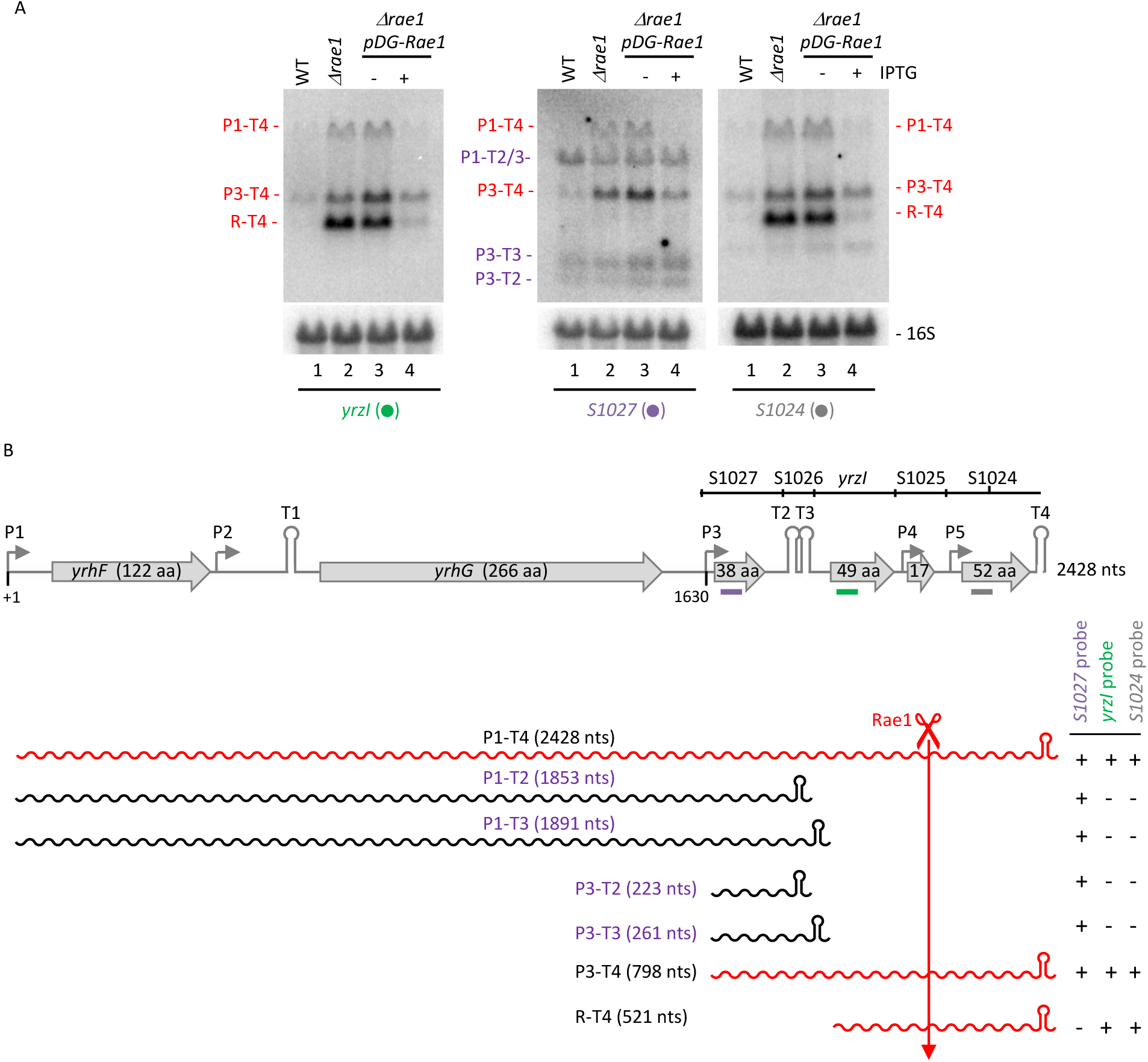
Mapping of the major *yrzI* operon transcripts. **(A)** Northern blots showing expression of the *yrzI* transcript in WT, *Δrae1* and plasmid complemented strains grown in the absence (−) and presence (+) of IPTG. Blots were probed with oligonucleotides complementary to *yrzI* (CC1589; green dot), *S1027* (CC1598; purple dot) and *S1024* (CC1600; grey dot) as indicated. Blots were rehybridized with a probe complementary to 16S rRNA (oligo CC058) as a loading control. **(B)** Structure of the *yrzI* operon and summary of transcripts identified in panel (A). ORFs are represented by grey arrows, the putative promoters by rightward-pointing arrows and putative transcription terminators by the hairpin structures. The sizes of the ORFs are given in amino acids (aa) and the lengths of the intergenic regions in nucleotides (nts) are indicated. The position of the 3 probes used is indicated by colors bars: purple for *S1027*, green for *yrzI* and grey for *S1024*. Transcripts from this locus are shown as wavy lines and those sensitive to Rae1 are in red. The presence or absence of the different species in the Northern blots shown in panel A is represented by (+) or (−) signs, respectively.

As seen previously [10], only the 2.4 kb (P1-T4), 0.8 kb (P3-T4) and 0.5 kb (R-T4) transcripts, which contain S1025, had Rae1-dependent expression profiles (Fig. 3A and 3B). The S1027 probe hybridized to the Rae1-sensitive P1-T4 and P3-T4 transcripts, as expected, and to three additional Rae1-insensitive species of ~1.9 kb, ~220 nts and ~260 nts (Fig. 3A, middle panel and Fig. 3B). These additional bands were not detected with the *yrzI* and *S1024* probes and their sizes are consistent with transcripts originating at P1 and P3 and terminating at T2 or T3, *i.e.* P1-T2, P1-T3 (indistinguishable in size on the Northern blot) and P3-T2 and P3-T3, respectively (Fig. 3A). The *S1024* probe hybridized to same three major transcripts as the *yrzI* probe (Fig. 3A, right panel). We also detected a very weak additional minor Rae1-independent species (around ~350 nts), whose origin remains unclear. We did not find any evidence for the use of the putative P2, P4 or P5 promoters under the rich medium conditions tested. More importantly, the most abundant (0.5 kb) species in the *rae1* deleted strain, R-T4, lacks an obvious promoter. We therefore concluded that it most likely results from processing of a primary transcript originating from P1 or P3.

### The 0.5 kb *yrzI*-S1024 transcript (R-T4) results from 5’ degradation by RNase J1 and is degraded via two alternative pathways

To determine the maturation event(s) that led to the production of the R-T4 species and which RNases eliminate the cleavage products, we performed high-resolution Northern blot analysis in different *B. subtilis* RNase mutants, containing or lacking Rae1.

In the absence of the 5’-exoribonuclease RNase J1 (*ΔrnjA*), the R-T4 band was replaced by a band migrating about 100 nts slower (referred as E1-T4), detected with both the *yrzI* and *S1024* probes (Fig. 4A, lanes 3 and 11). This suggests that the R-T4 species is generated by RNase J1 degradation from an upstream entry site, E1, until RNase J1 reaches the site labeled R (Fig. 4B). The E1-T4 species was more clearly visible in the *Δrae1 rnjA* double mutant strain, consistent with the ability of Rae1 to trigger its degradation (Fig. 4A, lanes 4 and 12). The fact that the R-T4 species was still present in a *Δrny rae1* strain (Fig. 4A, lane 8), would appear to suggest that the major endoribonuclease, RNase Y, is not responsible for cleavage at E1. However, we cannot rule out the possibility that, in the absence of cleavage by RNase Y, RNase J1 can gain access to the mRNA from a dephosphorylated 5’-end of the primary transcript to generate R-T4.

**Figure 4.**
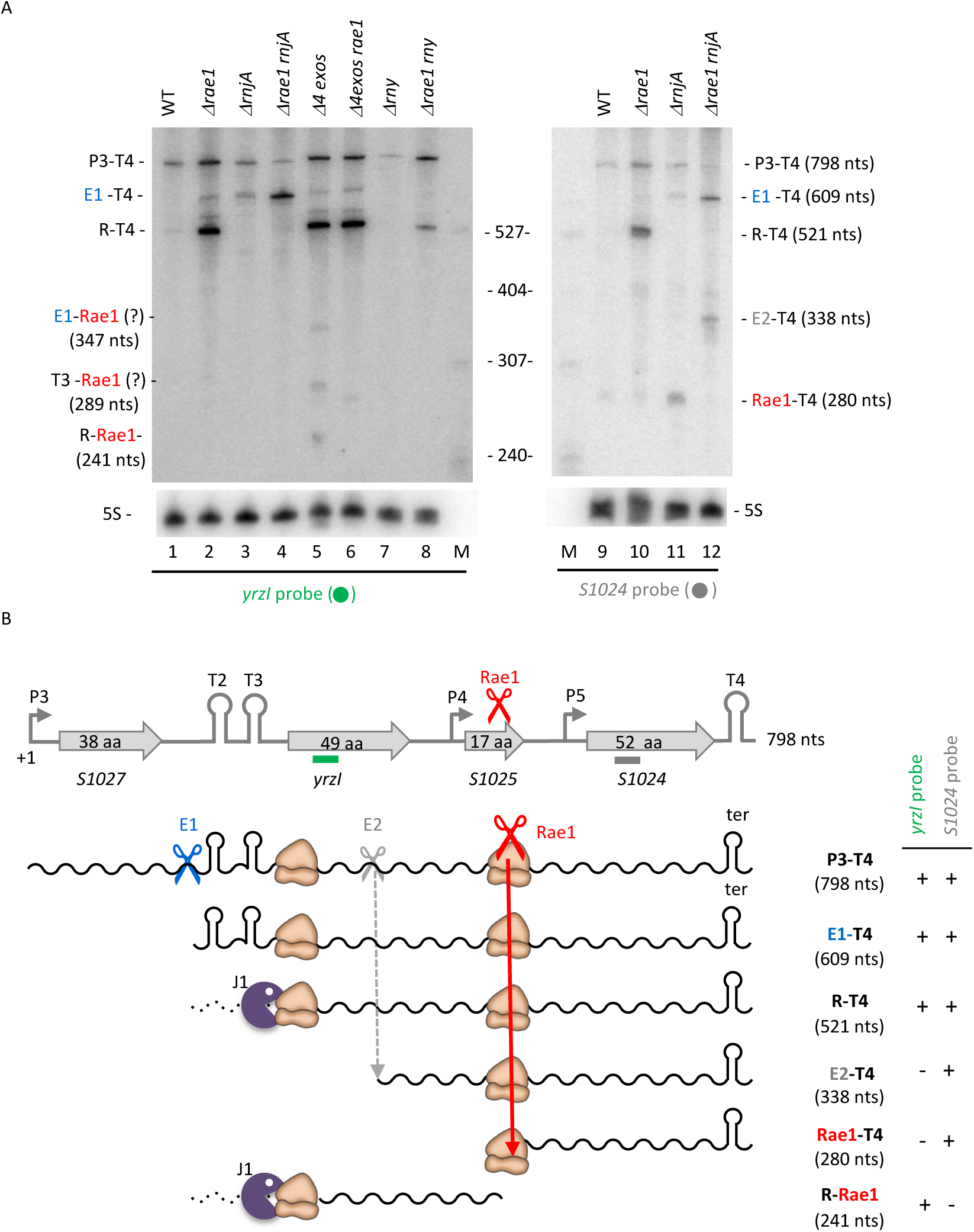
The 0.5 kb *yrzI*-S1024 transcript (R-T4) resulting from 5’-degradation by RNase J1 is degraded via two alternative pathways. **(A)** Expression of *yrzI* in different RNase mutants. High resolution Northern blot, probed with oligo CC1589 (*yrzI*; green dot) or CC1600 (*S1024*; grey dot), showing expression of *yrzI* in strains lacking specific *B. subtilis* RNases. Gene abbreviations are as follows: Rae1 (*rae1*), the four 3’-exoribonucleases PNPase, RNase R, RNase PH and YhaM *(4exos*), RNase J1 (*rnjA*), RNase Y (*rny*). Blots were rehybridized with a probe complementary to 5S rRNA (oligo HP246) as a loading control. **(B)** Summary of degradation pathways of short *yrzI* operon mRNAs. The structure of the region between S1027 and S1024 is shown. ORFs are represented by grey arrows, putative promoters by rightward-pointing arrows and transcription terminators by the hairpin structures. The sizes of the ORFs are given in amino acids (aa) and the positions of probes used are indicated by green and grey bars. The primary transcript (P3-T4) and the degradation intermediates are depicted. Endoribonucleases are symbolized by scissors and exoribonucleases by a Pacman symbol. Ribosomes protecting the 5’ end of the R-T4 transcript and involved in Rae1 cleavage are indicated.

Strikingly, the R-T4 transcript also strongly accumulated in a *Δ4 exos* strain, lacking the four known *B. subtilis* 3’-exoribonucleases (PNPase, RNase PH, RNase R and YhaM) but still expressing Rae1 (Fig. 4A, compare lane 5 to lane 1), suggesting that R-T4 can also be directly degraded from its 3’-end, independently of Rae1. This is unusual as 3’-terminator structures are not typically thought to be vulnerable to direct attack by 3’-exoribonucleases in *B. subtilis*. The R-T4, P3-T4 and E1-T4 species were each detected using a probe that hybridized to the T4 terminator confirming they all retain this structure at their 3’ ends (Fig. S1). Three shorter species were additionally detected in the *Δ4 exos* strain with the *yzrI* probe, all of which disappeared upon further deleting the *rae1* gene, indicating that they are all dependent on Rae1 cleavage and contain the S1025 ORF (Fig. 4A, lane 6). The shortest one likely corresponds to the upstream fragment resulting from Rae1 cleavage of R-T4 (labelled R-Rae1 in Fig. 4A, lane 5) consistent with the predicted size (241 nts) (Fig. 4B), while the two others could correspond to E1-Rae1 (347 nts) and T3-Rae1 (289 nts) respectively. In the *Δ4 exos rae1* strain, an additional degradation intermediate of ~270 nts was detected whose origin also remains unclear (Fig. 4A, lane 6).

The use of a probe complementary to S1024 permitted the detection of the ~280-nt downstream cleavage product of the R-T4 species by Rae1 (Rae1-T4) in the *ΔrnjA* mutant strain but not in the *Δrae1 rnjA* strain, as expected (Fig. 4A, compare lanes 11 and 12; Fig. 4B). The upstream and downstream fragments resulting from Rae1 cleavage were also detected in the *Δ4 exos* and *rnjA* strains, respectively, in global study to map sites of endoribonucleolytic cleavage to single nt resolution [19]. A longer species, which we call E2-T4 (~330 nts) (Fig. 4B), was also detected in the *Δrae1 rnjA* strain (Fig. 4A, lane 12). As for the R-T4 species, we confirmed that the E2-T4 species possessed the T4 terminator using the T4 probe (Fig. S1, lane 12). The 5’-end of the E2-T4 species corresponds to an additional entry site (E2) for RNase J1, previously mapped by primer extension assay to within the *yrzI* ORF [10]. The endoribonuclease responsible for the cleavage at E2 remains unknown.

Globally, these data indicate that the primary transcripts originating from P1 or P3 are cleaved by an unidentified endoribonuclease at the E1 site, followed by 5’-trimming by RNase J1 until it reaches the site labelled R to generate the stable intermediate R-T4. This intermediate can be degraded by either of two pathways: the first is initiated by Rae1 cleavage, followed by degradation of the products by 5’- and 3’-exoribonucleases, and the second is mediated by 3’-exoribonucleases from the 3’-extremity, independently of Rae1. A summary of the RNA processing/degradation pathways for the different RNAs of the *yrzI* operon, combining data from this and our previous paper [10] is shown in Fig. 4B.

### The R-T4 species is protected from RNase J1 degradation by ribosomes initiating translation of *yrzI*

The *yrzI* ORF has an exceptionally strong Shine-Dalgarno (SD) sequence, with 11 out of 12 possible Watson-Crick (WC) base pairs with the 3’ end of 16S rRNA (Fig. 5A). We therefore postulated that the 5’-end of the R-T4 transcript might result from the blocking of RNase J1 by ribosomes tightly bound to the *yrzI* SD sequence, similar to the phenomenon reported for the *hbs* and *cryIIIA* mRNAs that possess exceptionally strong SD, or SD-like sequences, respectively [20–23]. The hall-mark of these mRNAs that are protected from RNase J1 degradation is a ribosome ‘heel-print’ that occurs 8 nts upstream of the first G-residue of the consensus GGAGG SD sequence (Daou-Chabo et al., 2009) (Fig. 5A).

**Figure 5.**
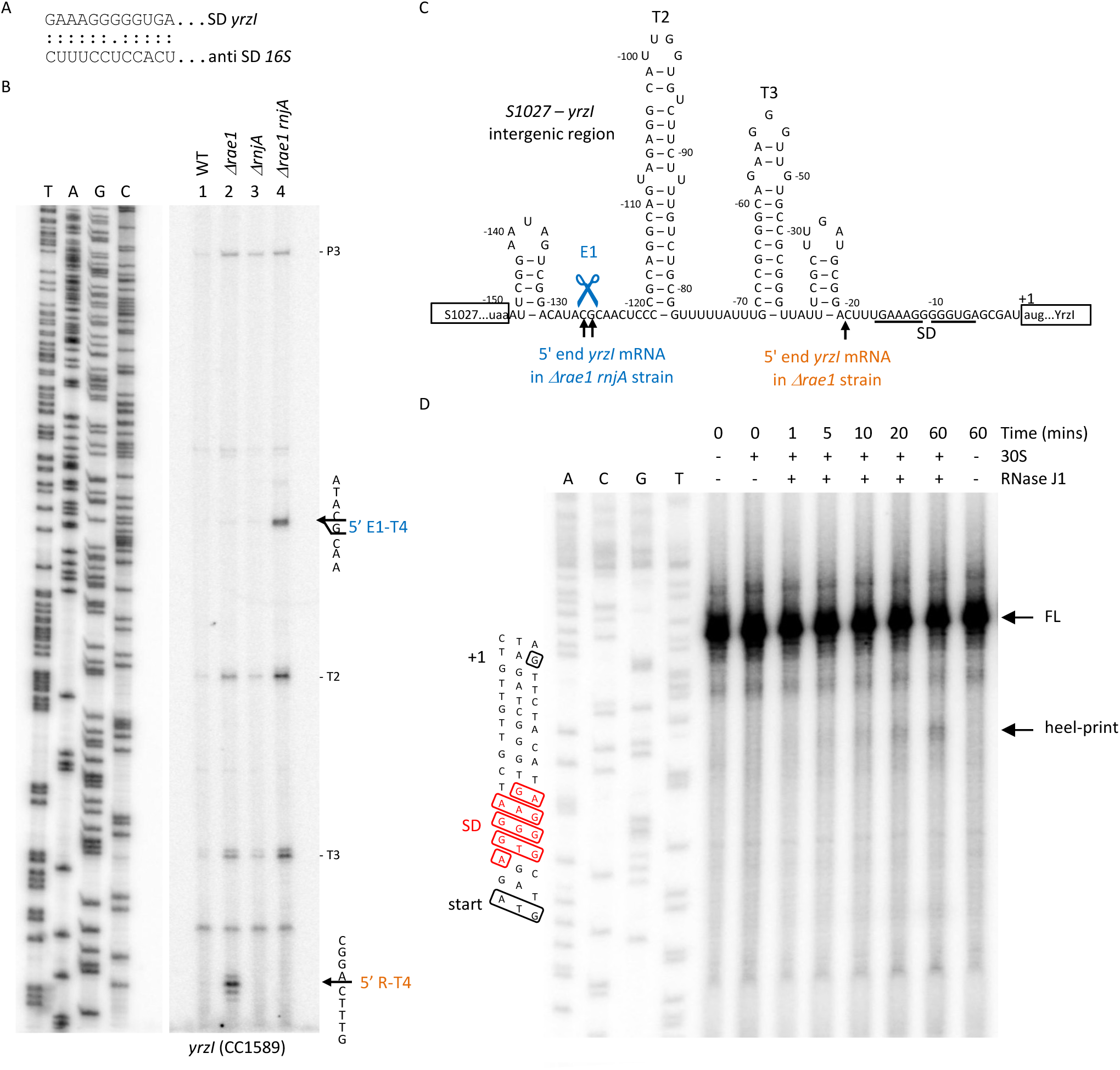
The 5’ end of the 521-nt (R-T4) transcript is protected by a ribosome bound to the *yrzI* SD sequence. **(A)** Potential base pairing between the *yrzI* SD sequence and the 3’ end of 16S rRNA. **(B)** Primer extension assay (oligo CC1589) on total RNA isolated from wild-type (WT) and strains lacking Rae1 (*Δrae1*), RNase J1 (*ΔrnjA*) and both (*Δrae1 rnjA*). Sequence lanes are labelled as their reverse complement to facilitate direct read-out. Mapped 5’ ends are shown to the right of the autoradiogram. Reverse transcriptase (RT) stops corresponding the T2 and T3 terminator structures and the predicted P3 promoter are also indicated. **(C)** Mapping of RNase J1 and Rae1-dependent 5’ ends to the predicted secondary structure (RNAfold; http://rna.tbi.univie.ac.at/cgi-bin/RNAWebSuite/RNAfold.cgi) of the S1027-*yrzI* intergenic region. Co-ordinates are given relative to the AUG start codon of YrzI (boxed). The predicted T2 and T3 terminator stem loops are shown. Nucleotides complementary to the anti-Shine Dalgarno sequence of *B. subtilis* 16S rRNA are underlined (SD). **(D)** Protection by 30S ribosomes against RNase J1 degradation of the *yrzI* transcript. The autoradiogram shows a heel-print assay on a 3’-labelled (p^32^Cp) RNA fragment of the *yrzI* transcript in the presence or absence of 30S ribosomes, incubated with RNase J1 for various times. A sequencing reaction on the template DNA was loaded on the left of the gel and labelled as reverse complement to facilitate direct read-out. Note: there is a slight difference in migration (equivalent to about 5 nts for the full-length (FL) transcript and about 4 nts for the heel-print) between the DNA sequence and the RNA reactions due to the 16 Da difference in MW between DNA and RNA nts.

To identify the precise 5’-ends of the main *yrzI* transcripts in *rnjA+ versus rnjA-*cells, we performed primer extension analysis on total RNA isolated from WT, *Δrae1, ΔrnjA* and *Δrae1 rnjA* double mutant strains, using a primer hybridizing to *yrzI*. In all four strains, we detected a 5’-end corresponding to the predicted P3 promoter upstream of S1027 (Fig. 5B). The calculated size of the fragment extending from P3 to the terminator following S1024 is 798 nts, corresponding well to the 0.8 kb fragment seen on Northern blots. In *Δrae1* cells, a 5’-end mapping to nt −21 relative to the *yrzI* start codon was strongly visible compared to WT cells (Fig. 5B). This 5’ end was located precisely 8 nts upstream of the G-rich motif in the *yrzI* SD sequence (Fig. 5A and C), identical what was observed for the *hbs* and *cryIIIA* ribosome heel-prints [22,23]. The predicted size of a transcript extending from this position to the transcription terminator downstream of *S1024* (T4) is 521 nts and correlates well with the size of the R-T4 band seen on Northern blots. In the *Δrae1 rnjA* double mutant, the 5’ end at nt −21 was no longer visible and was replaced by a species mapping to position −125/126, corresponding to the E1 site (Fig. 4A).

To show that the 521 nt *yrzI* species results from direct ribosome protection of the 521 nt *yrzI* species from RNase J1 activity, we transcribed a portion of the *yrzI* gene (−55 to +80 relative to the start codon) using T7 RNA polymerase. The transcript was 3’-labelled with ^32^P-pCp, pre-incubated with 30S ribosomal subunits and then subjected to RNase J1 degradation *in vitro* for different times. A protected species of the expected size accumulated over time in samples containing 30S subunits (Fig. 5D), confirming that the 5’ end of the 0.5 *yrzI* mRNA is protected from RNase J1 degradation by ribosomes initiating translation of the *yrzI* ORF.

Altogether, these data indicate that RNase J1 gains access to the *yrzI* transcripts at the upstream E1 site and degrades until it is blocked by a ribosome bound to the *yrzI* SD at nt −21, giving rise to the R-T4 species.

### Degradation of the *yrzI* polycistronic mRNA by 3’-5’ exoribonucleases

The data presented above showed that the highly stable R-T4 species can be degraded from its 3’ end independently of Rae1. We therefore analyzed the ability of individual 3’-exoribonuclease to promote R-T4 degradation by analyzing the profile of *yrzI* transcripts in strains deleted for one or more 3’-exoribonucleases (Fig. 6A). The R-T4 species was not detected in triple mutant *ΔyhaM rph rnr* that only retains PNPase (Fig. 6A, compare lane 7 with lane 5), but was observed in all strains lacking PNPase, albeit at a lower level than in the Δ*4 exos* strain (Fig. 6A, compare lanes 6-10 with lane 5). Thus, PNPase appears to be the major, but not the only 3-exoribonuclease involved in the degradation of R-T4. The R-Rae1 species was only detected in the *Δ4exo* mutants (Fig 6A, lane 5). Furthermore, a species migrating slightly faster than the upstream product of Rae1 cleavage (R-Rae1) was additionally observed in all strains lacking PNPase, including the triple mutant *Δpnp rph rnr* that only retains YhaM (Fig. 6A, lane 6, asterisk). This suggests that YhaM catalyzes the initial shortening of R-Rae1 and that PNPase is the principal enzyme that finishes the job of degrading it.

**Figure 6.**
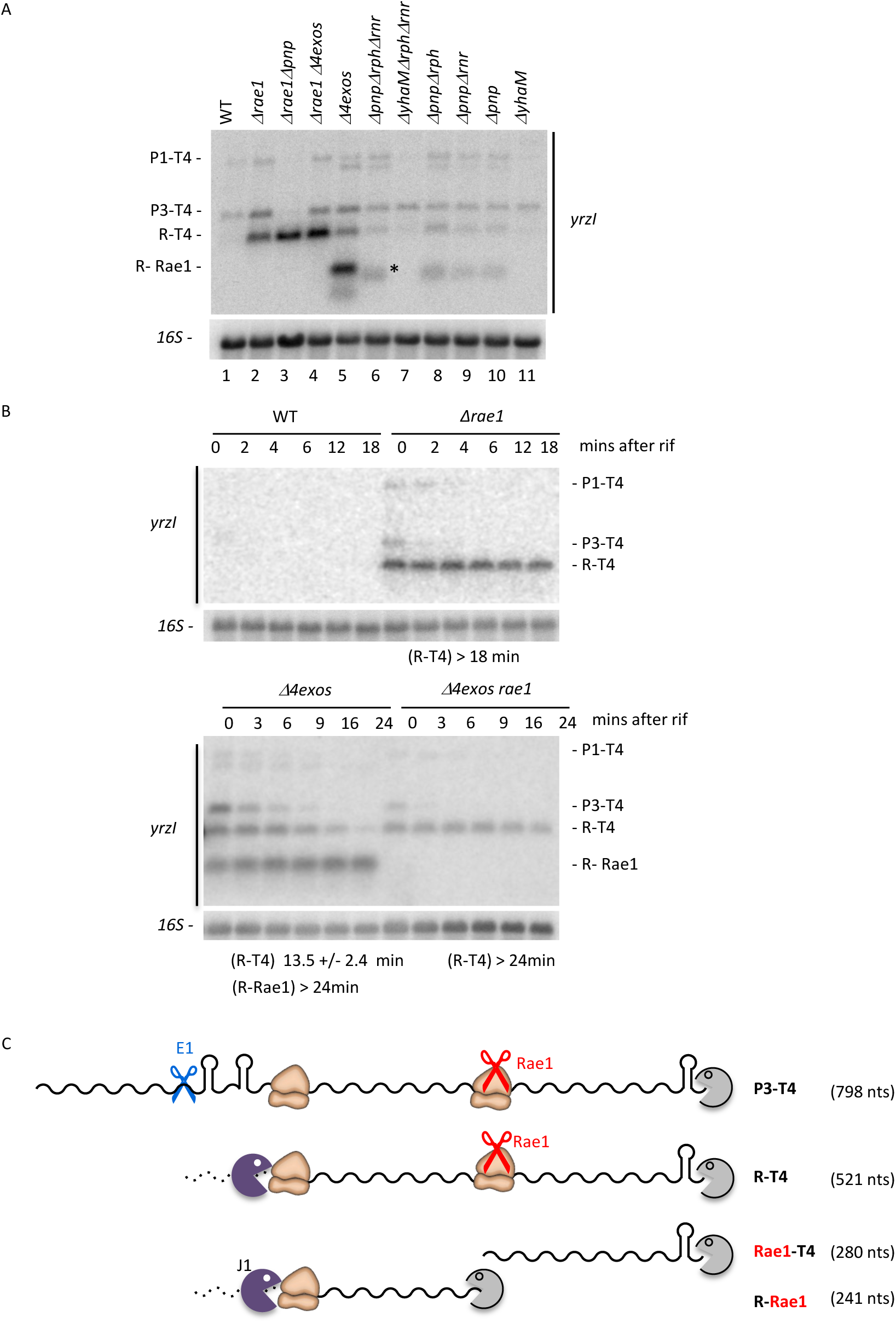
Degradation of the *yrzI* polycistronic mRNA by 3’-5’ exoribonucleases. **(A)** Northern blot, probed with oligo CC1589 (*yrzI*), showing expression of *yrzI* in strains lacking specific *B. subtilis* RNases. Gene abbreviations are as follows: Rae1 (*rae1*), the four 3’-exoribonucleases PNPase, RNase R, RNase PH and YhaM *(4exos*), PNPase (*pnp*), RNase PH (*rph*), RNase R (*rnr*) and YhaM (*yhaM*). **(B)** Half-lives were determined by Northern blot analysis in WT, *Δrae1, Δ4exo* and *Δrae1Δ4exos* strains at mid-log phase in rich medium at times after rifampicin addition, probed with oligo CC1589 (*yrzI*). Blots in panels A and B were rehybridized with a probe complementary to 16S rRNA (oligo CC058) as a loading control. **(C)** Degradation pathways of R-T4 transcript. The primary transcript (P3-T4) and the degradation intermediates are depicted. Endoribonucleases are symbolized by scissors and exoribonucleases by a Pacman symbol.

Lastly, we examined the relative contribution of the Rae1 cleavage pathway and the 3’-exoribonuclease pathway to R-T4 degradation by measuring its stability after addition of rifampicin to the culture medium in the *Δrae1* and *Δ4exo* mutants (Fig. 6B). The R-T4 transcript, which had a half-life of around 13.5 mins in the *Δ4 exos* strain, but >18 mins in either the *Δrae1* strain or the *Δrae1 Δ4exos* quintuple mutant, suggesting that the greater contribution comes from the Rae1 pathway (Fig. 6B). The fact that the half-life of the R-T4 transcript increases in the *Δ4 exos* strain compared to WT, despite the presence of Rae1, confirms that the effect of the quadruple mutant is at the post-transcriptional level and this 3’-degradation pathway can indeed attack the 3’ end of the mRNA independently of Rae1 (Fig. 6C). The co-existence of two independent rate-limiting degradation pathways could be reconciled if each pathway were to function on a different pool of mRNAs, *e.g* one that is translated (Rae1) and one that is untranslated (4 exos).

## Discussion

In this paper, we attribute functions to two of the four peptides encoded by the polycistronic *yrzI* mRNA from *B. subtilis:* the short 49-aa inhibitory peptide, called YrzI, and its antidote, a 17-aa peptide called S1025. We also describe a second naturally occurring case in *B. subtilis* of a ribosome bound to an exceptionally strong SD sequence protecting an mRNA fragment from 5’-degradation by RNase J1 (the other example being *hbs*) [21–23]. We further describe the non-canonical degradation pathways required to eliminate this species involving the translation-dependent endoribonuclease Rae1, on the one hand, and direct attack from the 3’-end by exoribonucleases on the other.

The YrzI peptide is highly conserved among the *Bacilli* (Fig. S2 A-B). A sequence encoding a peptide highly homologous to S1025 was identified downstream of the *yrzI* ORF in all analyzed Bacillus genomes, except for *B. cereus* (Fig. S2A and 2B). In *B. cereus*, YrzI peptides that exhibits lower homology to YrzI from *B. subtilis* are present in seven contiguous copies, but the ORF located immediately downstream does not share significant homology with the *B. subtilis* S1025 peptide (Fig. S2C). Although the *yrzI* operon is expressed most of the time in *B. subtilis,* pointing to a general role in the cell’s physiology, it shows peaks of expression in the cold, in stationary phase and late in sporulation [18], suggesting it may have specific functions under these conditions.

There are some intriguing parallels between the YrzI/S1025/Rae1 triad and classical toxin-antitoxin (TA) systems. As currently observed in TA modules, the YrzI toxin and S1025 antitoxin are encoded within the same operon. TA modules have been divided into seven classes according to the mode of action of the antitoxin [24–26]. We had previously identified a type II TA system in *B. subtilis* : the toxin EndoA (an RNase member of the MazF/PemK family of bacterial toxins) and the YdcD antitoxin protein encoded by the gene immediately upstream [27]. Our data suggest that YrzI/S1025/Rae1 triad could potentially constitute a toxin-double antitoxin system, with components resembling both the type II (protein antitoxin) and type V (RNase antitoxin) TA systems. On one hand, protection by S1025 against the toxic effect of YrzI was observed even in the absence of Rae1 indicating that the S1025 peptide is the major antidote to the YrzI toxin. On the other hand, Rae1 could easily counteract YrzI toxicity by promoting the degradation of the toxin-antitoxin mRNA, if S1025 peptide levels were insufficient to fully neutralize YrzI. Clearly, Rae1 does not serve as an antitoxin in cells growing in rich medium as a *Δrae1* strain has no major phenotype under these conditions. However, we cannot exclude that there may be certain growth conditions where expression of YrzI and S1025 become uncoupled and S1025 levels are insufficient to do the job alone. In this case the double protection afforded by Rae1 could be beneficial. A parallel can be drawn with the RatA/TxpA and YonT/as YonT type I TA systems of *B. subtilis* where the toxin-encoding mRNAs are degraded by RNase III [28]. However, in those cases inactivation of RNase III in rich medium is lethal.

Our data indicate that the *yrzI* and *bmrBCD* operons are both governed at the transcriptional level by AbrB and at the post-transcriptional level by Rae1 [12], suggesting there may be some as yet uncovered link between the two. Post-transcriptional modulation of *yrzI* expression first involves stabilization of the R-T4 fragment through cleavage by an unidentified endoribonuclease at the E1 site followed by 5’-timming by RNase J1 until it is blocked by a ribosome bound at the SD sequence of *yrzI*. The stabilized mRNA is then principally degraded through translation-dependent cleavage by Rae1. About 200 genes are predicted to have exceptionally strong SD sequences less than 20 nts upstream of their ORFs in *B. subtilis*, with maximum 2 mismatches out of 12 with the 3’-end of 16S rRNA equivalent to the experimentally confirmed *hbs* gene. Thus, at a conservative estimate at least 5% of *B. subtilis* genes are predicted to benefit from 5’-protection by ribosomes initiating translation at their SD sequences (Table S3). Note, however, that the original stabilizing motif in the *B. thuringiensis cryIIIA* mRNA (Stab SD) expressed in *B. subtilis* had only 8/12 consecutive base pairs [29] and so the actual number of mRNAs falling into this category may be much higher.

The *yrzI* polycistronic mRNA is targeted by several RNases: two endoribonucleolytic sites (E1 and E2) cleaved by unidentified RNases were mapped in addition to the Rae1 cleavage site. Remarkably, 3’-exoribonucleases can also participate in the degradation of the mature R-T4 transcript, despite the presence of the hairpin structure of the transcriptional terminator T4 at its 3’ end. Although Rho-independent transcription terminators are often described as providing resistance against 3’ degradation by exoribonucleases, the phenomenon has been understudied in bacteria and is likely to be dependent on the intrinsic stability of the RNA hairpins. Indeed, the predicted stability of the transcription terminator at the end of the *yrzI* operon is 11.1 kcal/mol, well below the average stability of experimentally determined Rho-independent terminators in *B. subtilis* of 16 kcal/mol [30]. This likely explains why the 3’ degradation pathway comes into play for this particular mRNA that is protected by ribosomes at the 5’ end. Although the R-T4 transcript was strongly stabilized in the *Δ4 exos* mutant, it showed no signs of decay over the time-course of the experiment in strains lacking Rae1. This unusual scenario, in which two rate-limiting degradation pathways target the same mRNA, can be explained if we consider that translated *yrzI* operon mRNAs are degraded by Rae1 and inaccessible to the 3’-exo pathway because of the flow of ribosomes to the end of the transcript, while untranslated *yrzI* mRNAs are eliminated principally by 3’-exoribonucleases.

This study of the *yrzI* polycistronic mRNA turnover highlights the complex interplay between ribosomes and the mRNA degradation machinery: degradation of the most abundant mRNA R-T4 species, generated by ribosomes obstructing 5’-exoribonucleolytic degradation, is influenced by the translational status of the mRNA for Rae-dependent cleavage and likely also for the 3’-exoribonuclease pathway.

## Acknowledgments

We thank lab members for helpful discussion. This work was supported by funds from the CNRS and Université Paris Cité (UMR8261), the Agence Nationale de la Recherche (ARNr-BasRae1, Labex Dynamo and Equipex Cacsice).

## Notes

### Competing Interest Statement

The authors have declared no competing interest.

