## supplemental Figures and Table for "Unconventional mRNA processing and degradation pathways for the polycistronic y*rzI* mRNA in *B. subtilis*"

### Supplementary Figure Legends

**Figure S1. Validation of presence of T4 terminator at 3' end of *yrzI* transcripts (A)** Reprobing of the high resolution Northern blot shown in Figure 4 with oligo CC3495 (complementary to terminator T4), showing expression of *yrzI* in strains lacking specific *B. subtilis* RNases. Gene abbreviations are as follows: Rae1 (*rae1*), the four 3'-exoribonucleases PNPase, RNase R, RNase PH and YhaM *(4exos*), RNase J1 (*rnjA*), RNase Y (*rny*). **(B)** Summary of degradation pathways of short *yrzI* operon mRNAs. The structure of the region between S1027 and S1024 is shown. ORFs are represented by grey arrows, putative promoters by rightward-pointing arrows and transcription terminators by the hairpin structures. The sizes of the ORFs are given in amino acids (aa) and the positions of probes used are indicated by green and black bars. The primary transcript (P3-T4) and the degradation intermediates are depicted. Endoribonucleases are symbolised by scissors and exoribonucleases by a Pacman symbol**.** Ribosomes protecting the 5' end of the R-T4 transcript and involved in Rae1 cleavage are indicated.

**Figure S2. Conservation of *yrzI* operon in Bacilli. (A)** Genomic island of *yrzI* among Bacilli using BioCyc.org [35]. **(B)** Conservation of YzrI and S1025 amino acid sequence in Bacilli. **(C)** YrzI and S1025 peptides in *Bacillus cereus*. Alignments were performed using Clustal Omega [36] in panels B and C. Conserved amino acid are marked with an asterisk.

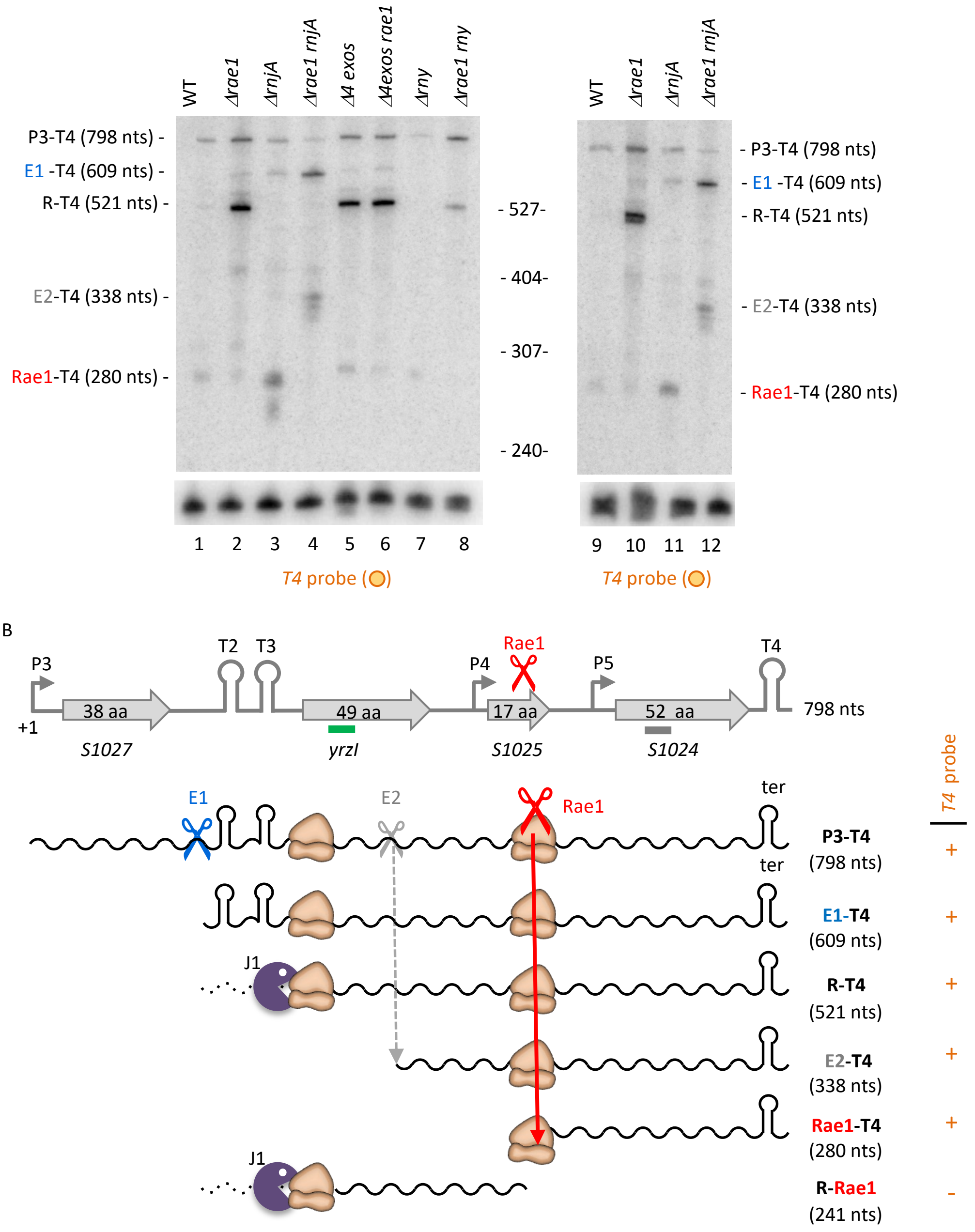


**Figure S1: Validation of presence of T4 terminator at 3' end of *yrzI* transcripts**

A

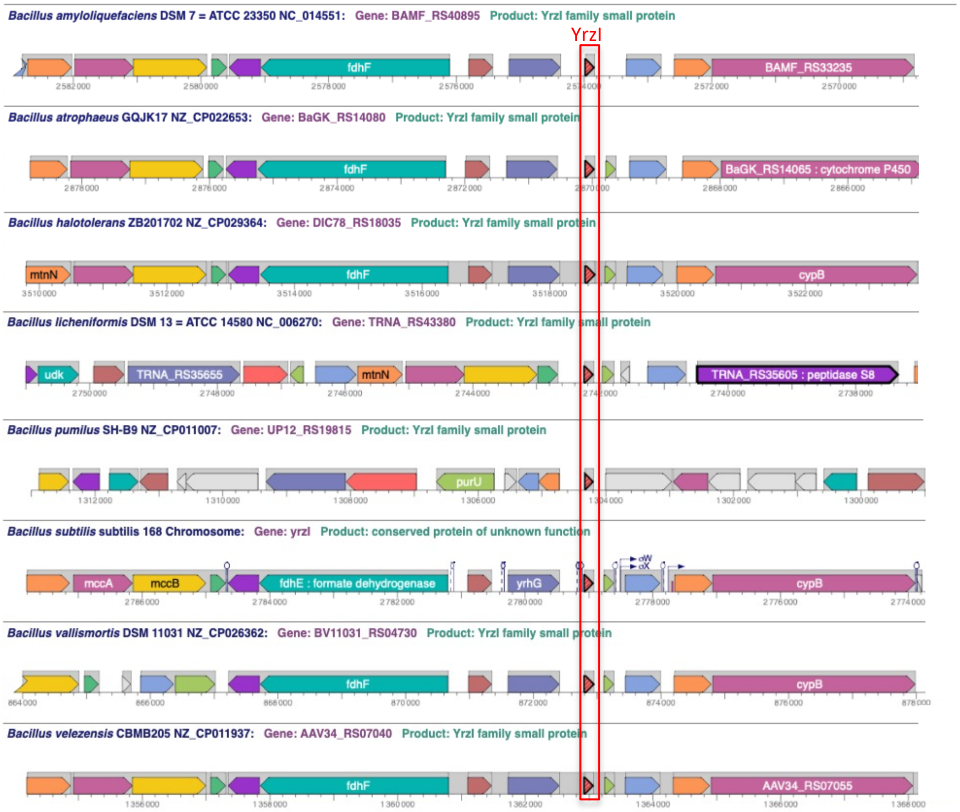


B

```
YrzI                                                                       aa    % ident
B subtilis              MTINLFFLTLTIKKRFKSLEEFEREQQIEQIYDEMKESQLKHLYLTNWR  49
B vallismortis          MTINLFFLTLTINKRFKSPEEFEREQQIEQIYGEMKESQLKHLYLTNWR  49     93,9
B atrophaeus            MTINLFFLTLTIKKRRKSLEEYEQEQQIERFYNETKETQLKHLYLTNWR  49     83,7
B amyloliquefaciens     MTINLFFLTLTISKRFKSAEEYEREQQVEEFYNEAKERQLKHFYMN---  46     73,9
B mojavensis            MTINLFFLTLTINKRFKSPEEFEREQQIEQIYDEMKESQLKHLYLTNWR  49     95,9
B velezensis            MTINLFFLTLTISKRFKSAEEYEREQQVEEFYNEAKERQLKHFYMN---  46     73,9
B licheniformis         MTINLFFLTLTIQRRFKSPEEYEREYEIEQMYNEMKDLQLKYLYLNSQR  49     71,5
B halotolerans          MTINLFFLTLTINKRFKSPEEFEQEQQIEQIYDEMKESQLKHLYLTNWR  49     93,8
B pumilus               MTIHLFFITLTIRRKYKDIETVLYEQQVETLYEEMKQKQLKYMN-----  44     53,3
Alignment               ***:***:****  :: *. *     *  ::* :* * *: ***::
S1025                                                                      aa   % ident
B subtilis              MLRMELLMHYTMEKDQV                                  17
B vallismortis          MLRMELLMHYTMEKDQV                                  17     100
B atrophaeus            MFRIELLIHYTMEKDQV                                  17     82,3
B amyloliquefaciens     MFRIELLNHYTMEKDQV                                  17     82,3
B mojavensis            MLKMELLMHYTMEKDQV                                  17     94,1
B velezensis            MFRIELLNHYTMEKDQV                                  17     82,3
B licheniformis        MIYRRELLAHYTREKDQV                                  18     70,6
B halotolerans          MLKMELLMHYTMEKDQV                                  17     94,1
B pumilus              MFMMLAILDCYTMEKDQV                                  18     52,9
Alignment               :    :*  ** *****
```

C

```
YrzI                                                                  aa    % ident
B. subtilis     MTINLFFLTLTIKKRFKSLEEFEREQQIEQIYDEMKESQL---KHLYLTNWR  49
B. cereus       MTFHIFFFTTVLQKNTLSETEINRKQQLKQLTDKITDIKSSYYTQMY-----  47     29,8
                MKFHLFFLTITIQKETISQNELKQERQYKQVIDEIRDRRSKYYTHL------  46     36,9
                MTISVLFLSITIQRNTISKDEILHNEQIEKAMNDVKERQALYCDHL------  46     32,6
                MKFKVFFLTITIQKNTFSESEMFHERQINQAMDHVKERQSHYCSHL------  46     39,1
                MKFKVFFLTITIQKNKFSESEMFHERQINQAIDHVKERQSHYCSHL------  46     39,1
                MKFKAFFLTITIQKRKLSQKEILREQHIQSIMDEVKERQSSYYTRLF-----  47     44,7
                MKVKAFFLTITIQKRKLSQKEISREQHIQNIMDEVKERQSSYYTRFF-----  47     44,7
Alignment       *..  :*:: .:::.  *  *: ::.: :.  .:: : :    ::
S1025                                                                 aa   % ident
B. subtilis      MLRMELL-MHYTMEKDQV                                   17
B. cereus       MMLIARCKEMIWIVQRDST                                   19     17,6
Alignment        **   :: * : :::*..
```

**Figure S2. Conservation of *yrzI* operon in Bacilli.**

**Table S1.** Oligonucleotides used in this study.

| Oligo | Gene | Sequence |
|---|---|---|
| HP246 | 5S | ATCGGCGCTGAAGAGCTTAACTTCC |
| CC058 | 16S | CTTCGTAGGGCAAATCGCCGTCCCC |
| CC1589 | *yrzI* | CTGCTGTTCTCGTTCGAATTCTTC |
| CC1598 | S1027 | GTGCGATAAGCATCGGTTAAGGAAGGACAC |
| CC1600 | S1024 | GGAATACATACAGCCTTTGCTGCTGCATTG |
| CC1731 | *yrzI* | tata<u>aagctt</u>GGACTTTGAAAGGGGGTGAGCGATATG |
| CC1732 | *yrzI* | ata<u>gtcgac</u>aaaaaggccatccgtcaggatggccCTTCAGTTTTTTTCGTTGCTTTTAGCGCC |
| CC1808 | S1025 | ata<u>aagctt</u>AAACTGAAGGGGGAATGGCATATGC |
| CC1809 | S1025 | tata<u>gtcgac</u>aaaaaggccatccgtcaggatggccCTTGACAGCTATTTCGCTA<br>AACTTGGTCCTTCTCC |
| CC3042 | *yrzI* | CTGCTGTTCTCGTTCGAATTCTTCC |
| CC3176 | *yrzI* | cagtaatacgactcactataGGGTTATTTCGCTTGATGCGGAC |
| CC3495 | *yrzI T4* | GTTTTCTGCCCTCTTTCTAATGCCAATG |

Restriction sites are underlined

Non-hybridizing sequences are in lower case.

**Table S2.** *Bacillus subtilis* strains used in this study

| Strain | Genotype | Reference |
|---|---|---|
| SSB1002 | W168 *trpC*+ | Lab strain |
| CCB329 | W168 *yhaM*::pm | This study |
| CCB375 | W168 *rae1*::pMUTIN | [10] |
| CCB376 | W168 *rae1*::pMUTIN ery *rph*::spc *rnr*::tc *yhaM*::pm *pnp*::kan | This study |
| CCB395 | W168 *pnp*::kan | This study |
| CCB396 | W168 *rph*::spc *rnr*::tc *yhaM*::pm *pnp*::kan | [31,37] |
| CCB406 | W168 *rph*::spc *pnp*::kan | [38] |
| CCB407 | W168 *rph*::spc *rnr*::tc *pnp*::kan | This study |
| CCB409 | W168 *rnr*::tc *pnp*::kan | This study |
| CCB434 | W168 *rnjA*::spc | [39] |
| CCB441 | W168 *rny*::spc | [39] |
| CCB604 | W168 *rae1*::pMUTIN + pDG-*yacP* | [10] |
| CCB748 | W168 *rae1*::pMUTIN ery *rnjA*::spc | [10] |
| CCB761 | W168 *rae1*::pMUTIN ery *rny*::spc | This study |
| CCB815 | W168 *rae1*::pMUTIN + pDG-*yrzI-ter* | This study |
| CCB839 | W168 + pDGYrzI | This study |
| CCB913 | W168 + pDGYrzI-S1025 | This study |
| CCB914 | W168 *rae1*::pMUTIN + pDGYrzI-S1025 | This study |
| CCB915 | W168 + pDGS1025 | This study |
| CCB916 | W168 *rae1*::pMUTIN + pDGS1025 | This study |
| CCB1210 | W168 *rph*::spc *rnr*::tc *yhaM*::pm | This study |
| CCB1058 | W168 *abrB::*cm | [12] |
| CCB1059 | W168 *rae1::pMUTIN* ery *abrB::*cm | [12] |
| SSB1030 | W168 *pnp*::cm | [38] |

Table S3

Predicted exceptionally strong SD sequences (gaaaggaggtga or with one or 2 mismatches) in B. subtilis lying no more than 20 nts upstream of the start codon.
Only those with a central GGAGG or GGGGG motif were retained. The position in genome, the localization of the gene on strand (+) or (-), the distance of the SD sequence from the AUG codon, the number of mismatches and the name of the gene are indicated.

| pattern_position | pattern_strand | bp_from_start | mismatches | gene | product | pattern |
|---|---|---|---|---|---|---|
| 4550 | + | -16 | 1 | yaaB | hypothetical protein | TGGTGCGTTAGTGAAGTGAA-GAAATGAGGTGA-GCAATTGTATATTCATTTAG |
| 55847 | + | -18 | 2 | spoVG | septation protein SpoVG | ACTAATGCTTTTATATAGGG-AAAAGGTGGTGA-ACTACTGTGGAAGTTACTGA |
| 87617 | + | -16 | 1 | dusB | tRNA-dihydrouridine synthase | GCCAGAAAAATAGAACAAAT-GAAAGGAGGAGA-AAAATTGTTCAAAATCGGAG |
| 108657 | + | -16 | 2 | yacL | PIN and TRAM-domain containing protein YacL | TTGTGAAATTAAAATGTTAA-AAAAGGAGGTGG-GGGTATGTTAAAACGAATAG |
| 136350 | + | -18 | 2 | rplD | 50S ribosomal protein L4 | AATCTAAATAATTCTCTTAG-GAAAGGAGGAAA-TTGATTATGCCAAAAGTAGC |
| 140432 | + | -18 | 2 | rplN | 50S ribosomal protein L14 | AAGTTCGGAATTCTTTTTCC-GAAGGGAGGTAA-CAAATAATGATTCAACAAGA |
| 141954 | + | -19 | 2 | rpsH | 30S ribosomal protein S8 | GCTGGTAATCCCCAAGAAAG-GGAAGGAGGTAA-TTATATAATGGTTATGACAG |
| 142384 | + | -17 | 2 | rplF | 50S ribosomal protein L6 | TTTGGTAAGAAGTTTAACAT-GAATGGAGGTGT-TTGGTATGTCTCGTGTAGGT |
| 149934 | + | -18 | 2 | rplQ | 50S ribosomal protein L17 | ATTCAGAAACAGCATTCCAA-TAAAGGAGGGGA-CATCACATGTCATACAGAAA |
| 154282 | + | -17 | 2 | rpsI | 30S ribosomal protein S9 | ACGAACTTCGCGGTTAATTA-AAAAGGAGGTTA-CCAATTTGGCACAGGTTCAA |
| 159075 | - | -15 | 2 | gerD | spore germination protein GerD | TAAAGGTAAGACAAGTATGT-GAAAGGAGCTTA-AGCATGTCAAAAGCCAAAAC |
| 181359 | - | -17 | 2 | feuC | iron-uptake system permease protein FeuC | TTCCTTTATTTAATTAAACG-AAAAGGAGGGGA-GCAAAATGGCTAAAAAATAT |
| 264172 | - | -16 | 2 | glnT | sodium/glutamine symporter GlnT | TCAGCATTTTTTAAACATGA-AAAAAGAGGTGA-TTCTATGCAGCAAATCCTAG |
| 277143 | + | -16 | 2 | rtpA | tryptophan RNA-binding attenuator protein inhibitory protein | GTGGCATAGCAGATGAGTTA-GAAAGGAGGATA-TCAAATGGTCATTGCAACTG |
| 314868 | + | -14 | 2 | yceG | hypothetical protein | ATCGTTTCAATGACAAACCG-GAAAGGAGGATA-TGATGAGACAAGAAGAACTA |
| 322252 | + | -18 | 2 | opuAB | glycine betaine transport system permease protein OpuAB | ACAAACACAAGATCCTTCTG-CACAGGAGGTGA-AATAAGATGGATAGACTGCC |
| 402369 | + | -18 | 1 | srfAD | surfactin synthase thioesterase subunit | GGTTTCATAAATGAAGTGAT-GAAAGGAGGAGA-CAGCCAATGAGCCAACTCTT |
| 459849 | + | -17 | 2 | kipI | kinase A inhibitor | TCATCCTGATGAACGAGAAT-TAAAAGAGGTGA-GAAAGATGACTGTACGATAT |
| 471690 | + | -18 | 2 | ydaD | general stress protein 39 | ACAACAAATAGACATGAAAA-CAAAGGAGGAGA-ATCATCATGGCAAATTATCC |
| 515691 | + | -18 | 2 | acpS | holo-[acyl-carrier-protein] synthase | CCATTTAAATAGTACGTACG-CAAAGGAGGTTA-TCATACATGATTTACGGCAT |
| 545574 | + | -20 | 2 | yddJ | hypothetical protein | TTGTTTATACTAAATATTGA-AAAAGGAGGTAA-TGTTTATTTTGAAAAACCTT |
| 589702 | + | -14 | 2 | ydfJ | hypothetical protein | ATGTGATCAAGCAATGTGTT-GAAAGGAGATTA-TCATGTCAAAAATGTTATAT |
| 640643 | + | -18 | 2 | thiL | thiamine-monophosphate kinase | AAATAGGCATAAGCAGGGTT-GAAAGGAGGTTT-TGGTTCATGGATGAATTTGA |
| 647741 | + | -18 | 2 | tatAY | Sec-independent protein translocase protein TatAy | TAGAGGAAATCGAATAGAGG-GAAAGGAGGAGC-CCAAATATGCCGATCGGTCC |
| 682365 | - | -15 | 2 | ydjP | AB hydrolase superfamily protein YdjP | GCCATACTCATGATGGAAAC-TAAAGGAGGAGA-AACATGCCATACATCATATT |
| 686567 | - | -15 | 2 | gabP | GABA permease | TTCAGTAATGATTGCATTTT-AAAAGGAGGTAA-CTTATGAACCAGTCTCAATC |
| 754461 | + | -18 | 2 | yesE | hypothetical protein | GACTGAATAGTCTAGAAATA-AAAATGAGGTGA-TGATGAATGTTGATGAATGA |
| 755890 | + | -16 | 2 | cotJA | spore coat associated protein CotJA | ATTAAGGTCATTGTTTTTGA-GAAAGGAGGAGC-CGGAATGAAGGATATGCAGC |
| 760435 | + | -16 | 2 | yesN | transcriptional regulator | TGTGATCACAATACCGTGCC-GAAATGAGGTGG-TGTGATGTATAAAATACTGC |
| 800211 | + | -20 | 1 | yfnF | hypothetical protein | AATTCTTCAAAAAGTAATGT-GAAACGAGGTGA-ACAAGACTGTGACCAGCGCG |
| 804550 | - | -16 | 2 | yfnC | MFS transporter | GGTATGCTTTGGTTATATAT-GTAAGGAGGAGA-CTCAATGGCTATCGCCGCCC |
| 825720 | - | -17 | 2 | yfmD | Fe(3+)-citrate import system permease protein YfmD | CGTTTTTGCAACCAGACGCA-GTAAGGAAGTGA-GACTGTTGTATCATTCAGCC |
| 844236 | + | -16 | 2 | yflB | hypothetical protein | ATAAAATATAAGGTATAGAT-AAAAAGAGGTGA-AATGATGGATTTTTCCCATA |
| 847261 | - | -15 | 2 | yfkT | spore germination protein YfkT | ATTCCGTTTTGAACATGACA-GAAAGGATGTGC-TTTATGGACAAGACCTCTGC |
| 859506 | - | -16 | 2 | yfkN | nucleotide phosphoesterase | GCTGTAACACAAATTTCGTT-GAAAGGTGGAGA-ACTGGTGAGAATACAGAAAA |
| 876405 | + | -20 | 2 | dusC | tRNA-dihydrouridine synthase 2 | GTTCAGCCGTATCCCATAAT-GAAAGGATTTGA-TTTTTGTTATGACAGAAAAT |
| 882246 | + | -19 | 2 | acoL | dihydrolipoyl dehydrogenase | CGCAGCATTAATTTTATAGG-AAAAGCAGGTGA-AAACGACATGACATTAGCCA |
| 895599 | + | -19 | 2 | yfiC | ABC transporter ATP-binding protein | CATTTACGAGTCACAATTCG-GAAGGGAGGGGA-GCGAGTCATGCTAAAAGACA |
| 900059 | + | -20 | 2 | yfiG | metabolite transporter | TCGGGTGCTCTGCCATTCTT-GAAAGAAGGGGA-AGAAAGCCGTGAGTACAAAA |
| 912585 | - | -16 | 2 | yfiR | TetR family transcriptional regulator | TTATAGGTAAGAAAATTGCA-GAAAGCAGGCGA-TATCTTGTCACCAAAAGTAA |
| 916018 | - | -17 | 2 | yfiU | MFS transporter | TAATATATTTCAAGTGAACT-AAAAGGAGTTGA-CGAAAATGTCTGAAGCATTA |
| 926868 | + | -17 | 2 | yfhI | MFS transporter | AACTCGTTGTTTTTTAAAAT-CAAAGGAGGCGA-TGACGTTGAGTATCAAAAAC |
| 929389 | + | -16 | 1 | yfhL | hypothetical protein | CTTTCTGCACGTAACAATGA-GAAAGGAGATGA-TGATATGACGGGTTTAGTCG |
| 935443 | - | -15 | 2 | yfhP | hypothetical protein | TGAATGTGCAGGATCACCCT-GCAAGGAGGTCA-CCTATGGATACGGGCACACA |
| 938713 | + | -17 | 2 | ygaC | hypothetical protein | ACTAACTGTAGGGCTTTATA-GAAAGTAGGGGA-GAAATATGGGCTATCCCAAG |
| 976549 | + | -19 | 2 | yhbI | MarR family transcriptional regulator | ATCTTGTGAACTATTATTGA-GGAATGAGGTGA-AAAGGAGTTGACTGAATCAG |
| 977756 | + | -18 | 2 | yhcA | MFS transporter | CAGATCAGTCATCAAACCAT-GAAAGGAGGTTG-TCTTAGATGGCTACAGGCAA |
| 1024846 | + | -18 | 1 | yhdH | sodium-dependent transporter YhdH | ATAAGAGATAATATAGGCTG-GAAAGGAAGTGA-CGTTTATTGTCTGAGCAAAA |
| 1038636 | + | -16 | 2 | yhdX | hypothetical protein | ATGATCCTTTATAATAGCAA-AAAAGGAGGCGA-GATCATGGGAAAAGGGAGAA |
| 1062574 | + | -16 | 2 | natA | ABC transporter ATP-binding protein | ATTAGTATATAATGGGTCAG-GGAAGGAGGAGA-TTCAGTGCTTACAATTGATC |
| 1079405 | + | -16 | 2 | ecsC | ABC transporter substrate-binding protein EcsC | AGCACGGAAAGCTGTGAACT-GAAAAGAGGTAA-TCAAATGACAGATAATCAGC |
| 1133388 | - | -16 | 2 | yhjO | MFS transporter | CTACTACTATATCAAACAGC-GAAAGGCGGCGA-TCACATGTTGACCAGTTTTC |
| 1136142 | - | -18 | 2 | yhjR | hypothetical protein | TACAGAAGCCCAATTCCTAA-GAATGGAGTTGA-ATCCCCTTGCATTACAGCTA |
| 1149327 | - | -18 | 1 | gerPD | spore germination protein GerPD | GAGCATATTCCATCAGAAAT-GAAAGGAGATGA-GCAAGCATGATCTTTACAGT |
| 1205145 | + | -19 | 2 | yjaU | hypothetical protein | GATAGCGTGAGATGAAGTGT-GAAAGGAGGCGG-GCAAGCGATGGGAATAACGG |
| 1206609 | + | -19 | 2 | med | transcriptional activator protein med | AATACATGTTATAGCTTGTC-AAAAGGAGTTGA-ACATGACTTGATCACAAGGC |

| | | | | | | |
|---|---|---|---|---|---|---|
| 1240125 | - | -16 | 2 | yjbP | Bis(5'-nucleosyl)-tetraphosphatase PrpE | CATAATAAATAAAACAATCG-GTAAGGAGGAGA-CCGCATGGCTTACGATATCA |
| 1253084 | + | -18 | 2 | yjcZ | hypothetical protein | CTCACATAGTGTATAGAGAG-TAAAGGAGGTGC-TTATCTATGGGCTTTGGATA |
| 1297708 | + | -17 | 2 | yjlB | hypothetical protein | CGCTATATGTGGAAGACAAA-AAAGGGAGGTGA-AATTGATGGAAAGAGAGACA |
| 1300433 | + | -16 | 2 | uxaC | uronate isomerase | TTAACATTTTGAAATAGAAT-GAAAGACGGTGA-GGACATGGAACCCTTCATGG |
| 1304424 | + | -17 | 2 | yjmD | zinc-type alcohol dehydrogenase | GGAGCGTGTATGAATTCTTA-AAAAGCAGGTGA-GTAAAATGAAAGCGGTTCAA |
| 1330492 | + | -19 | 2 | ykzL | hypothetical protein | ATTAACGTAAAAGAATAATA-GAAACGAGGTGG-TCAGCTCATGCTCATTGAAC |
| 1345612 | + | -18 | 2 | xkdX | phage-like element PBSX protein XkdX | AAGCTATCATTACTTTCCTT-GAAAGGAGATTA-TGCTGAATGAATTATTGGGT |
| 1376836 | + | -18 | 2 | ykkD | multidrug resistance protein | ACACAGGATGAGACAGAGGA-AAAAGGAGGCGA-GGCATAATGCTGCACTGGAT |
| 1385612 | - | -16 | 2 | metE | 5-methyltetrahydropteroyltriglutamate--homocysteine methyltra | GAAAGAGAGTGTTTTACATA-TAAAGGAGGAGA-AACAATGACAACCATCAAAA |
| 1395353 | + | -17 | 2 | ykzD | hypothetical protein | CTTTTGGACAATCTAAGTAG-AAAAGGAGGCGA-AACGCATGGAAAAGAAAGAG |
| 1401585 | - | -18 | 2 | ykoQ | metallophosphoesterase | GATCACTTACTATACTGATA-GAAGGGAGGGGA-TATATTTTGTTTACAATCTT |
| 1413999 | - | -17 | 2 | sspD | small acid-soluble spore protein D | GTTAAGAATTTCTTTATCGA-AAAAGGAGATGA-AAAAGATGGCGAGCAGAAAT |
| 1505395 | + | -20 | 2 | yknY | ABC transporter ATP-binding protein | ATCTGATCAGGTGACTGACG-GAATGGAAGTGA-AATCCTAAATGATTCAGCTT |
| 1514035 | + | -16 | 2 | ykpB | oxidoreductase | TTTTTTTGATACCTAAATGT-GAAAGGAGATCA-CAACATGAAATTTTTGGTTG |
| 1565831 | + | -17 | 2 | ylbB | hypothetical protein | TGACGACAATAGCACTAGAC-TAAAGGAGGTTA-AACGGATGACAAAAATAAAA |
| 1581577 | + | -19 | 2 | ftsL | cell division protein FtsL | CGAAACGCGAACGCAAAATT-AAAAGGAGGTCA-TCAGCCTATGAGCAATTTAG |
| 1595916 | + | -18 | 2 | sbp | small basic protein | CTGAAACTGCTTAAAGACTG-AAAAGGAGGAGA-ATTGTCATGTGGCTGCCCGT |
| 1608902 | + | -16 | 2 | ylmC | hypothetical protein | TACTTGAACAAAAGGACGGA-AAAAGGACGTGA-TGGAATGATCAGCATTTCAG |
| 1621353 | + | -20 | 2 | pyrC | dihydroorotase | CGTGCCTTACAAACCAATGT-GAAAAGAGGAGA-AGCAGCGTATGTCATATCTC |
| 1623717 | + | -18 | 2 | carB | carbamoyl-phosphate synthase pyrimidine-specific large chain | GAAATGATCGAAACAACAGA-GAAAGAAGGGGA-AGCGGTATGCCAAAACGCGT |
| 1630363 | + | -18 | 2 | cysH | phosphoadenosine phosphosulfate reductase | GCTACTGTTTGTTTGTCTCC-GAAAGGAGGAAA-GAAGAAATGTTAACGTATGA |
| 1650365 | + | -18 | 2 | prpC | protein phosphatase PrpC | GCGCGAAGGAGCGTCAAGAC-GAGACGAGGTGA-TGAGGGTTGTTAACAGCCTT |
| 1665318 | + | -18 | 2 | acpP | acyl carrier protein | ACGAATCCACTATAATACTT-GAGGGGAGGTGA-ATTGCTATGGCAGACACATT |
| 1682562 | + | -17 | 2 | dprA | protein smf | CTGTTCCTGCTTTTTCTAAC-AAAAGGAGGTCA-ATCTATTGGATCAGGCCGCT |
| 1688658 | + | -17 | 2 | hslU | ATP-dependent protease ATPase subunit ClpY | ATACTGGAAGAGCTTGAATA-GAAAGGACTTGA-GGCGCATGGAAAAAAAACCG |
| 1692111 | + | -18 | 1 | fliE | flagellar hook-basal body complex protein FliE | GATGAAGGCGTTAGAAATCG-GAAAGTAGGTGA-ATGAATGTGATTAATGCAAT |
| 1737814 | + | -19 | 2 | ribC | riboflavin biosynthesis protein RibC | ATGCAAAAAAGCGAACAATA-GAAAAAAGGTGA-CCGTTCTGTGAAGACGATAC |
| 1767293 | + | -16 | 2 | ymdA | ribonuclease Y | GCAACAACCAAGTTCATAGC-AAGAGGAGGTGA-AAGTATGACCCCAATTATGA |
| 1781790 | - | -17 | 2 | ymcC | hypothetical protein | TTAAATAATACAACTGTATT-GAAAAGAGGGGA-ATGGATTGAACGGTATCGCA |
| 1863433 | + | -14 | 2 | ymaC | hypothetical protein | ATGAACCGCGCCATTCTTAT-GAAAGGAGTTAA-ACATGAGAAGATTTTTACTA |
| 1873585 | - | -18 | 2 | cwlC | sporulation-specific N-acetylmuramoyl-L-alanine amidase | GCCGTGTTTTTTCATATCCT-GTAATGAGGTGA-TGAAAAATGGTTAAAATTTT |
| 2007404 | - | -18 | 2 | ftsR | LysR family transcriptional regulator | CAATATCACTTTTCATCATA-GCAAGCAGGTGA-GAGCGCGTGGAAAGCGGAGA |
| 2024029 | - | -17 | 2 | yoaC | sugar kinase YoaC | CACTGAAAACAGGGGGATAA-GAAAGGAGGCTA-ACAGAATGAAAAAACAAAAA |
| 2037372 | - | -16 | 2 | yozS | hypothetical protein | TCTACCAGTTTGATTGGCTT-TAAAGGAGGAGA-AGAAATGAAAGCACTCATAT |
| 2083017 | - | -16 | 2 | yobU | transcriptional regulator | TCTTTGTCTATAATCTTAAA-AAAAGGAGGAGA-TCGTATGGGCTTTTCACATA |
| 2097622 | - | -17 | 2 | yocK | general stress protein 16O | GCAAAACGGGAAACAGGATA-GAAAGGAGAGGA-TATACATGGCACTGACAAAA |
| 2098842 | + | -16 | 2 | yozN | hypothetical protein | ACCGCCAGCATACAACTATA-GAAAGGAGGTTT-GCCTGTGGTTCAACCAACTC |
| 2099110 | + | -16 | 2 | yocN | hypothetical protein | AAGCGGTTAACTCCTCAATC-AAAAGGAAGTGA-TGAGATGTTCTTTTCACCCT |
| 2120748 | + | -18 | 2 | rsbRC | RsbT co-antagonist protein RsbRC | ATTCGCAAAAACAACACATT-TAAAAGAGGTGA-TCAATCATGGCAAAAAACAA |
| 2124003 | + | -17 | 1 | gerT | spore germination protein GerT | TATACTATTTTGTCCGTGAA-GAAAGGATGTGA-AGACAATGTTTGAGTGGAAC |
| 2134225 | + | -18 | 2 | yodI | hypothetical protein | ATATGCTGATAGAAAGTCAG-GATAGGAGGGGA-ACAATATTGGAGAGATATTA |
| 2156124 | - | -16 | 2 | yotB | hypothetical protein | AGAAATAAATTTTTCATTAA-GAAAGGTGGAGA-CAAAATGAAAATTGATTATG |
| 2157114 | - | -16 | 2 | yosX | hypothetical protein | ACTAATATTTTATTCAAACT-TTAAGGAGGTGA-TTCATTGGAAGTAGGAGACA |
| 2161782 | - | -16 | 2 | yosP | - | CGGAAAAAAATATTTCAATT-GAAAGGACATGA-TTAATTGACAAAAATTTATG |
| 2161088 | - | -14 | 1 | yosQ | HNH homing endonuclease YosQ | GCAGCGAAGCTATGCGAACC-CAAAGGAGGTGA-AAATGATAAGGAAAGAAGTC |
| 2161782 | - | -16 | 2 | nrdF | ribonucleoside-diphosphate reductase subunit beta | CGGAAAAAAATATTTCAATT-GAAAGGACATGA-TTAATTGACAAAAATTTATG |
| 2168875 | - | -16 | 2 | yosD | hypothetical protein | TTCAATTTTTAAAGAGACTT-ATAAGGAGGTGA-ATAAGTGGGACGCCATAAAG |
| 2169790 | + | -16 | 2 | yosA | hypothetical protein | ATACCATATACTATCCTTAC-TAAAGGAGGTGG-CAATATGGGATTCTACAGCG |
| 2186816 | - | -17 | 2 | yorE | hypothetical protein | ATAATAAAATCGAGTTAAAG-GGAAGGAGGCGA-ATTACTTGGAGGTACTTGGG |
| 2193666 | - | -17 | 2 | yoqU | hypothetical protein | AATAAAATATTTATTTTAAT-GGAAGGAGGTAA-CCGTAGTGCCAAATCAAGAG |
| 2206583 | - | -20 | 2 | yopQ | hypothetical protein | AACTTTAAATTTAGCAACAT-GACATGAGGTGA-AAGGCTATATGACAGTGATC |
| 2207763 | - | -17 | 1 | yopP | integrase/recombinase YopP | ATTGATGAGTTATTAGATGA-GAAAGGAGGAGA-AGATAATGCAAAATAAAATT |
| 2208984 | - | -16 | 1 | yopL | hypothetical protein | TTTTAGAATTACTTTTACTT-TAAAGGAGGTGA-GACAATGAAAAAACTTATTA |
| 2210158 | - | -16 | 2 | yopK | hypothetical protein | ACCAATAAAGACAACAAAAA-GAAATGAGGGGA-AATAATGGAGTTAATAAGGA |
| 2219965 | - | -17 | 1 | yonT | hypothetical protein | ATGTTATAATAGAGTCATAG-GAAAGGAGGTGT-ACATAGTGCTTGAGAAAATG |
| 2222323 | + | -16 | 2 | yonP | hypothetical protein | TGTATATAATAAATCCATAA-GTGAGGAGGTGA-GATAATGCAAAAGGATTCAG |
| 2227484 | + | -20 | 2 | yonJ | hypothetical protein | CTTCCTGTTAAAGGCGTTGA-GTAAGGAGATGA-CTGACTGAATGATCGATCCT |
| 2232666 | + | -15 | 1 | yonE | hypothetical protein | GAATTAGTCTATTTTTAAAA-GAAATGAGGTGA-AACATGGTAACCTTAAATAA |
| 2234214 | + | -18 | 1 | yonD | hypothetical protein | CTATTTAAAACTTCACTTTT-GAAGGGAGGTGA-AATTATTTGACAAAGAAGCA |
| 2239064 | + | -18 | 2 | yomW | hypothetical protein | ATAAAATTGGTTGCCAAAAA-GAAAGTTGGTGA-AAAGACATGAGCATGACTGT |
| 2249134 | + | -19 | 1 | yomI | transglycosylase YomI | TCTTTGTCCTCTCCCCTACT-GAAAGGAAGTGA-TTCTTACTTGAGTCAAAACC |
| 2259456 | + | -18 | 2 | yomF | hypothetical protein | GGGAAGCCTACCAACAAAAT-TAACGGAGGTGA-ACTAAACTGGCTGATTTTGC |

| | | | | | | |
|---|---|---|---|---|---|---|
| 2265674 | - | -15 | 2 | bdbB | disulfide bond formation protein B | CACAATATTGAACACGTCCT-GAAAGGAATTGA-AGTATGAATACAAGATATGT |
| 2275701 | - | -16 | 2 | yokJ | hypothetical protein | ATTTAATCAATACAACTTTA-GAAAGTAGGTGC-GGACTTGAGCATTGACATGT |
| 2280865 | + | -15 | 2 | yokE | hypothetical protein | CAACTATGCTTGTGAACGAT-GAAAGGAGATAA-GTGATGGGACAAAAATTTCA |
| 2281649 | + | -17 | 2 | yokD | AAC(3) family N-acetyltransferase | GTATCTAATAAGAGGTTATA-GATAGGAGATGA-ACTAAATGAAAAAAATAGTT |
| 2310404 | + | -14 | 2 | ypdQ | ribonuclease H-like protein | AAAAAATGATACCTAATGAT-GAAAGGAGTTCA-TAATGCCTACAGAAATATAT |
| 2318057 | - | -16 | 2 | bcsA | chalcone synthase | ATTGTTGCGGCACAATGTAT-GCAAAGAGGTGA-TCGCATGGCGTTTATTTTAT |
| 2332080 | - | -17 | 2 | ypsB | cell cycle protein GpsB | AAATATATGTGATGATTCAC-GATACGAGGTGA-AAAAGATGCTTGCTGATAAA |
| 2371502 | - | -17 | 2 | hisC | histidinol-phosphate aminotransferase | TTTAGCGGCTTGTACAGTTT-AAAATGAGGTGA-CTGATTTGCGTATCAAAGAA |
| 2385826 | - | -17 | 2 | hbs | DNA-binding protein HU 1 | CGAACTGAATGTATCCTTTT-GGGAGGAGGTGA-AAGGCATGAACAAAACAGAA |
| 2413478 | - | -17 | 2 | aroD | 3-dehydroquinase | AATAAAACAAGGTAAAGATT-GAAAGGATTTGA-GACGAGTGAACGTGTTAACG |
| 2443310 | - | -18 | 2 | spoVAA | stage V sporulation protein AA | ATACTACATATATAACCACC-GAAAGATGGTGA-TCAATGATGGAACGACGAAT |
| 2455780 | - | -18 | 2 | aspA | aspartate ammonia-lyase | AAAACGATAACCAGCAAAGA-GAAAGAAGGTTA-AAGATTATGTTAAACGGCCA |
| 2483885 | + | -18 | 2 | yqjG | membrane protein insertase MisCB | TTTTATAAACCGCATTTATA-AAAAGGAGGAGA-ACAAAATTGTTAAAAACATA |
| 2504667 | - | -15 | 2 | ptb | phosphate butyryltransferase | AAAAGGCGGAGTCGAAACAA-GAAAGGTGGTAA-CAGATGAAGCTGAAAGATTT |
| 2509643 | - | -16 | 2 | prpD | 2-methylcitrate dehydratase | GCGATATCAAACAAAATCAT-GAAAGGAGCTGG-TAGAATGCCGAAAACGGATC |
| 2516423 | + | -16 | 2 | yqiG | NADH-dependent flavin oxidoreductase YqiG | GAATATGTGCGAATCCAAAC-GAAAGAAGATGA-TCAAATGAATCCTAAGTATA |
| 2535529 | - | -16 | 2 | spoIIIAE | stage III sporulation protein AE | ATACCTTCTATGTCATAACC-GAAAGGAGGCGG-TAGATTGAAGCGCTTTCAAT |
| 2537618 | - | -17 | 2 | spoIIIAA | stage III sporulation protein AA | TTTAAAGAAGCCAATCGGTA-GAAAGGAGGAGG-CTCTGTTGAATGAAATCGCT |
| 2594179 | - | -17 | 2 | yqfS | endonuclease IV | AAGAAACCAATCTAAGAAAA-GAAAGTAGGGGA-ACCTGTTGCTGAGAATAGGC |
| 2595652 | + | -16 | 2 | yqfQ | hypothetical protein | TTCTAATAGCAGAGAGAGCT-GAAAGGAGGTAC-AAGTATGTTTTCACCGCAGC |
| 2598629 | - | -18 | 2 | yqfO | hypothetical protein | CGCCGATCGAATGGAGCTGT-TAAAGGAGGTAA-TCGATCATGGCTAAAAGTGT |
| 2618450 | - | -18 | 2 | yqfA | hypothetical protein | TCGTGAGAGAAATTTAAATA-GAAACGAGGAGA-AGTTATATGGATCCGTCAAC |
| 2630862 | - | -17 | 2 | hemN | oxygen-independent coproporphyrinogen-III oxidase 1 | TGTACTGCGGCTTCTTTCTG-TAAAGAAGGTGA-ATCCGTTGAAATCAGCTTAT |
| 2641196 | + | -17 | 2 | comER | ComE operon protein 4 | TGGTTTAGAGGATAATAGCT-CAAAGGAGGGGA-AACCATTGAAGATAGGCTTT |
| 2651618 | + | -13 | 2 | yqeB | hypothetical protein | GGAATTCGATTTTTTCAGTA-AAAAGAAGGTGA-CATGTTGCAGAATCAATCTC |
| 2661087 | + | -14 | 2 | yqcG | hypothetical protein | TTTTGTTATGACGATTAAAT-GAAAGGATATGA-TCATGAAAGTATTTGAAGCC |
| 2672045 | - | -16 | 2 | yqbQ | hypothetical protein | CAAAAATTAAAAATACCGCA-GTAAGCAGGTGA-TGACATGATAGAACTTTTCG |
| 2677469 | - | -18 | 2 | yqbO | hypothetical protein | CTGTTAGAAAAAGAAGCGAA-GAAGGGAGGTAA-TTAACTATGGCTAAACTAAC |
| 2678223 | + | -16 | 0 | yqdB | hypothetical protein | TTAGCTTATCCTCCGAATAT-GAAAGGAGGTGA-AATTATGTCGACCTATGAAT |
| 2680993 | - | -17 | 2 | yqbK | hypothetical protein | GTTGATTTCTCAGTTTCTTC-AAAAGGAGGTCA-AATAGATGAACGGCGGAACT |
| 2685134 | - | -17 | 2 | yqbD | hypothetical protein | ATTAGAAGTCTTTTCTTATT-TTAAGGAGGTGA-ATAACATGCCAAGAGAATTG |
| 2688305 | - | -15 | 2 | yqbA | hypothetical protein | GACCGCAAAGATCAAGACCA-GGAAGGAGGTAA-AGCATGTCAAAAAAATCTGT |
| 2690850 | - | -17 | 2 | yqaR | hypothetical protein | CTCTTTTGCCGATAATTAGT-AAAAGGAGGGGA-TGAAATTGATTCAAATAGAT |
| 2710792 | - | -17 | 2 | yrkJ | hypothetical protein | GAAAGATACCCTTTTCCTAT-GAAAGGAGGCAA-ATAAAATGGATATTGCCTTT |
| 2721657 | - | -17 | 2 | yrdR | transporter | TTCTCATTGTAATTTTAAAG-AAAAGGAGGGGA-ATTCAATGAAAAAAGGTCAA |
| 2752782 | + | -19 | 2 | yraG | spore coat protein F | TCGGGAAATGTAACGTTGAT-AAAAGGAGGTTA-CATCAATATGGATCATCAAA |
| 2779077 | - | -17 | 1 | yrzI | hypothetical protein | TTTCGCTTGATGCGGACTTT-GAAAGGGGGTGA-GCGATATGACGATTAACCTA |
| 2794132 | - | -16 | 2 | yrrO | protease YrrO | AAGGAAACGGTTTATTAATC-AAAAGGAGGTTA-AGCTATGACTGCCGTAAATG |
| 2805484 | - | -17 | 2 | yrzQ | hypothetical protein | CAGCATCAATTTTTAATCAT-GAAAGGAGATGC-GCAGGATGAATCGGACGATG |
| 2818479 | + | -14 | 2 | yrvJ | N-acetylmuramoyl-L-alanine amidase YrvJ | TGCTTTTTTAATGATCATCA-GAAAGGAGGACA-CAATGAACAAGAAATACTTT |
| 2840092 | + | -17 | 2 | yrzF | serine/threonine protein kinase | AACAGCTGCATAAAATAGAG-CAAAGAAGGTGA-ATGAAATGCTGACAAAAGCA |
| 2843832 | - | -16 | 2 | yrbC | transcriptional regulator | AAAGTGTACATAGAAAATGT-AAAAGAAGGTGA-AAACATGGCAGGCCATTCCA |
| 2855169 | - | -17 | 2 | rpmA | 50S ribosomal protein L27 | GCGTGTGACCACAAACATAA-TATAGGAGGTGA-GCTACATGCTTAGATTAGAT |
| 2872754 | - | -17 | 2 | spoVID | stage VI sporulation protein D | TCTAATTTTAAGGATTAGAT-GAAAGGAGGATA-TGAACTTGCCGCAAAATCAT |
| 2874177 | - | -17 | 2 | hemL | glutamate-1-semialdehyde 2%2C1-aminomutase | CGGAGTAATTTTATTCAGTT-GACAGGAGTTGA-GGCAGATGAGAAGCTATGAA |
| 2898913 | + | -17 | 2 | ysnF | stress response protein YsnF | AACGATATCATGCAAATTCA-TAAAGGAGGAGA-TTGATATGAAAAGCATAGTT |
| 2925901 | - | -16 | 2 | yshA | cell division protein ZapA | TAAGAACTAGGATTCTCGCG-GAATGGAGGAGA-AACGTTGTCTGACGGCAAAA |
| 2973164 | - | -17 | 2 | mutM | formamidopyrimidine-DNA glycosylase | TACGATGCGAAATAAACAGA-GATAGGAAGTGA-TGGATGTGCCGGAATTACCA |
| 2988714 | - | -18 | 2 | accA | acetyl-CoA carboxylase carboxyltransferase subunit alpha | AATCTGCTGGATATGCATCA-AACAGGAGGTGA-CATTGAGTGGCTCCAAGATT |
| 3000971 | - | -16 | 2 | ytnL | hydrolase YtnL | AGAAAATATATTGGGATAGG-GAAAGGAGCGGA-ACATATGTCTTTGGATTATT |
| 3021051 | - | -16 | 2 | sppA | signal peptide peptidase SppA | TATAATGAGAGAGTTTAGGA-AAAAGGAGGAGA-AAAAATGAATGCAAAAAGAT |
| 3025659 | - | -17 | 2 | sspA | small acid-soluble spore protein A | TTTTGACACATTCTATACTC-ACAAGGAGGTGA-GACACATGGCTAACAATAAC |
| 3056324 | - | -16 | 2 | ytoP | aminopeptidase YtoP | TAAAATAAAAGGACTACATA-GAAATGAGGGGA-AAACATGAATCAAGAAACGA |
| 3083232 | - | -19 | 2 | yteP | multiple-sugar transport system permease YteP | CAGTGAATGCAAGAATCCGT-GAAAGGAGGCTA-GGAAAAAATGAAAACAGCAG |
| 3092194 | - | -19 | 2 | bioD | ATP-dependent dethiobiotin synthetase BioD | GCAAACATTTCATTCAATCG-GAAAGGAGCTGC-ACATCATTTGAGGGGTTTTT |
| 3134125 | + | -18 | 2 | ytlD | ABC transporter permease | GACACTGTTTCAAACGATAT-GGAAGGAGCTGA-ATTCTCTTGAGAAACAGCAA |
| 3145024 | - | -17 | 2 | mntB | manganese transport system ATP-binding protein MntB | TACTAAAGCGCTTAAATAAA-GAAAAGAGGTGG-AGGATATGTTCCCTGTTGAG |
| 3149720 | - | -16 | 2 | menB | 1%2C4-dihydroxy-2-naphthoyl-CoA synthase | GACTCATTCACATAGAGAAT-AAAAGGAGGTCA-TCATATGGCTGAATGGAAAA |
| 3153989 | + | -17 | 2 | yteA | hypothetical protein | CTAAAGGTATGCATCACAGA-GAAACGAGGCGA-TCACATTGCTTACGAAAGAA |
| 3246371 | - | -16 | 2 | yufS | hypothetical protein | CAATCCTGATCAAGCTGAAG-GAAAGGAGCTGG-TCATGTGAAAACAAAAGTCG |
| 3251704 | + | -19 | 2 | mrpF | Na(+)/H(+) antiporter subunit F | GGAGTCATTTGAAAAAGCCA-TACAGGAGGTGA-GCCGGTGATGTTTACGCTGA |
| 3292300 | - | -16 | 2 | dhbA | 2%2C3-dihydro-2%2C3-dihydroxybenzoate dehydrogenase | TGTTTGGCATTTAGAGCGAA-GAAAGGAATTGA-TGATATGAATGCAAAGGGTA |

| | | | | | | |
|---|---|---|---|---|---|---|
| 3312741 | - | -17 | 2 | yuxL | peptidase YuxL | AGAAAAACGAAATACATGTA-AAAAGGAGGAGA-TGAATATGAAAAAGCTGATA |
| 3319565 | - | -16 | 2 | yutD | hypothetical protein | ATTATGATAGGAATGAGTGA-AAAAAGAGGTGA-GATCATGATTCTTATTCAAA |
| 3321438 | + | -16 | 2 | lytH | L-Ala--D-Glu endopeptidase | AAACTATGTGTATTCAATAT-GAAAGGAGATTA-TTTTGTGAAAGTTTTGTTAT |
| 3326771 | - | -18 | 2 | yunF | hypothetical protein | AAAACACGTGTGATAAACTG-GAAGAGAGGTGA-ATGAGTGTGATTTATATCGG |
| 3335397 | + | -16 | 1 | yuzJ | hypothetical protein | ATATCTTATCCTATCTCCAC-TAAAGGAGGTGA-CATTATGGGATTCTACAATT |
| 3336276 | - | -16 | 2 | pucE | xanthine dehydrogenase subunit E | GTTGCTTGAAGCGATCGACA-GAAAGGGGCTGA-AGCAATGGACATAAAAGAGG |
| 3348826 | - | -17 | 2 | frlM | ABC transporter permease | AATGAAGTTCTTTAAAACCG-GAAAGGAGGAGT-AACTGATGCTGCCTCAGAAG |
| 3349710 | - | -19 | 2 | frlN | ABC transporter permease | CTTCGCACCTTTTCTTTTTA-TAAAGGAGATGA-GATACCCTTGGTCAATCAGA |
| 3385705 | + | -18 | 1 | cssR | transcriptional regulatory protein CssR | AACTAGTTTATAATGACGTT-GAAAGGATGTGA-AGAGCCTTGTCATACACCAT |
| 3390764 | + | -17 | 2 | gerAA | spore germination protein A1 | TAACCTCTACTAAGGTTTTG-GATAAGAGGTGA-CCTCATTGGAACAAACAGAG |
| 3414045 | - | -18 | 2 | yvrO | ABC transporter ATP-binding protein | TGATCCCGCGGTCAAGGAAG-GAAAGCAGGTTA-CAGTCAATGCTGACACTGAA |
| 3421610 | - | -17 | 2 | sspJ | small acid-soluble spore protein J | TTAAACATGTACCTATTTTT-TAAAAGAGGTGA-TACGAATGGGTTTCTTTAAT |
| 3468240 | - | -17 | 2 | opuCD | glycine betaine/carnitine/choline transport system permease p | AACATCATTATTTTGACTAA-GAAAAGAGGTGG-ATCATATGGAAGTACTACAG |
| 3517494 | - | -18 | 2 | epsL | sugar transferase EpsL | TTAAAACGGTTTCGAAAAAC-GAAAGGAGCTGT-GAATCTTTGATCCTGAAACG |
| 3570527 | - | -16 | 2 | whiA | sporulation transcription regulator WhiA | TACTGAAAGAATGAAGCCTT-GAAATGAGGTGG-CTATATGTCATTTGCATCAG |
| 3643563 | - | -17 | 2 | comFA | ComF operon protein 1 | TCATTCAGGCATACTGTTTC-GAAAGGAGGCGT-GCTATGTGAATGTGCCAGTT |
| 3714682 | - | -15 | 2 | ywrJ | hypothetical protein | AAGTGATGCCCTTTTTCAGT-TAAAGGAGGAGA-ATCGTGAAAGGTTTGAATCA |
| 3720907 | + | -17 | 2 | ywrC | HTH-type transcriptional regulator YwrC | AATGATTGCTAAAAGGCTAC-GAAAGCAGGTGG-AGCAATTGAGTCATGAATAT |
| 3723437 | + | -16 | 2 | ywqM | HTH-type transcriptional regulator YwqM | CAATCGCTAAAACCGATTTA-GAAAGGAAGAGA-ACAAGTGGAATTGAAGCAGC |
| 3743663 | - | -16 | 2 | mscL | large-conductance mechanosensitive channel | TTTTTTCTTTACAAATATAG-AAAAGCAGGTGA-TTGTATGTGGAATGAATTTA |
| 3759595 | - | -16 | 2 | ywnJ | hypothetical protein | ACATGTAGGCACGTACGTTA-TAAAGGAGGTTA-TGTCATGAACCGCCTATTGC |
| 3775333 | - | -16 | 2 | ywmD | hypothetical protein | TAGTTGGAAAATATGAAAAA-GAAAGGAAGTAA-TCATATGAAAAAATTGCTGG |
| 3780544 | + | -17 | 2 | ywmA | hypothetical protein | TTACATGGTATACTTTATTG-GAAAGGAGCTGG-CAAATGTGAATCATTCTTTT |
| 3783806 | - | -17 | 2 | atpG | ATP synthase gamma chain | CTTTTTACGCAGATGAAGAG-AAAAGGTGGTGA-AATCTTTGGCCTCATTACGC |
| 3787619 | - | -19 | 2 | atpB | ATP synthase subunit a | AGCTTAAACGTTCATCAATG-GAAGAGAGGTGA-AAACCCTTTGAATCATGGTT |
| 3789063 | - | -20 | 2 | upp | uracil phosphoribosyltransferase | AATTGTTGATGGGACAATAA-AAAAGGAGCTGA-AACACAGTATGGGAAAGGTT |
| 3819746 | - | -14 | 2 | ywjB | hypothetical protein | ATTCGGGCAAATGGTGCATC-GATAGGAGGAGA-ACATGGAAAGAAAAACTGTG |
| 3844381 | - | -16 | 2 | ywhL | hypothetical protein | GCTGCATGTTGTGCATTTCT-AAAAGGACGTGA-AATCATGAAAAAGAATGATC |
| 3845775 | - | -16 | 2 | ywhK | hypothetical protein | GCGTGAGCAGTGTATGTCTG-AAAAGGAAGTGA-AGGAATGAGAAAAAAACAAGC |
| 3888866 | - | -16 | 2 | spsE | spore coat polysaccharide biosynthesis protein SpsE | CTATTATCATATTTGGCCGG-GCAAGGAGGCGA-AATAATGGCAGCGTTTCAGA |
| 3899810 | - | -17 | 2 | ywdE | hypothetical protein | ATATACATAGAATTTCTATG-TAAAGGAGGTGT-ATGTCGTGAGGATTAGCTCT |
| 3926401 | - | -14 | 2 | ywbO | hypothetical protein | GTTTAAAATCAATTTAGTAT-GAAAGGAGATTA-TCATGACAGTACACATTAAA |
| 3929112 | - | -16 | 2 | efeM | iron uptake system component EfeM | GGCAAAACATTTTACAAACG-GAAAGGAACTGA-CAAAATGAATTTCACAAAAA |
| 3935575 | + | -15 | 2 | ywbE | hypothetical protein | AATGACTATAATGAACAAAA-GAAAGGATGTGC-GTTATGAACGGGCAAGAGCG |
| 3938232 | - | -14 | 2 | ywbB | hypothetical protein | GAAAGGAATGTTAGAATATG-TAAAGGAGGTGC-GCATGCTAGATTATATATGG |
| 3939853 | + | -15 | 2 | epr | minor extracellular protease epr | TATATATCAAACATCATAGA-AAAAGGAGATGA-ATCATGAAAAACATGTCTTG |
| 3950589 | - | -17 | 2 | ywaC | GTP pyrophosphokinase YwaC | GACAGCGGATAAAGTTCCGT-TAAAGGAGATGA-CGAACATGGATTTATCTGTA |
| 3960178 | - | -17 | 1 | licA | PTS system-lichenan-specific transporter subunit IIA | CAAAGAAGCGGCTCTGTAGA-GAAAGGAAGTGA-ATCGCGTGAATGAGGAAATG |
| 3984029 | - | -15 | 2 | yxkH | polysaccharide deacetylase | AAAGGAGGAAGAAATAGACA-GAAAGCAGGAGA-ACGATGAAACGATTGTTTTT |
| 3992654 | + | -16 | 2 | yxjN | hypothetical protein | TTTTCGCTATGCTTAATGCA-GAAAGGATTTGA-TGACATGGCGATGAGCCCTT |
| 3995056 | + | -18 | 2 | pepT | peptidase T | CCCGCCATGCTAAAATAAGA-CAAAGGAGATGA-TTGGAAATGAAAGAAGAAAT |
| 3997946 | + | -17 | 2 | yxjH | hypothetical protein | TTTACATGCTTAATATTTCA-GAAAAGAGGCGA-ATAACATGGCTCAACAAACG |
| 4114120 | + | -20 | 2 | gntK | gluconokinase | CTCTCACAACAAATGCTGGC-AAAAGGAGCTGA-ATACAACAATGACTAGTTAT |
| 4139541 | - | -16 | 2 | yycO | hypothetical protein | TACCTTTCCTACACTCACAA-GAAAAGAGGAGA-TGTAATGAAGTTGAAGAAAC |
| 4141340 | + | -17 | 2 | phrG | phosphatase RapG inhibitor | GATTGAAGAACTGATTAAAC-GAACGGAGGTTA-TATAAATGAAAAGATTTCTG |
| 4166592 | - | -16 | 2 | yybS | hypothetical protein | TCTATTCGTATGTTCAGGAT-GATACGAGGTGA-CTAAGTGAAACAAACGAGAG |
| 4189389 | + | -16 | 2 | yyaO | hypothetical protein | ACCTGAATATAACCAACTTA-GGAAGAAGGTGA-TCCCATGCCAAATAAAAGTA |
| 4191181 | + | -16 | 2 | yyaL | hypothetical protein | TTAAATATACACTAATTATA-GGAAGAAGGTGA-TCCCATGCCAAACAAAAGTA |
| 4198848 | - | -18 | 2 | rpsR | 30S ribosomal protein S18 | TTATCGCCTAAAATGAAAAA-GAAAGGAGGGAA-ATGACAATGGCAGGAGGACG |
| 4199738 | - | -18 | 2 | rpsF | 30S ribosomal protein S6 | CATTATGGGCCGCTTAGTCC-AAAAGGAGGTGC-AAACAGATGAGAAAGTACGA |
| 4203336 | - | -18 | 1 | yyzM | hypothetical protein | CAATAGATACAGCAGTTTAG-GAAAAGAGGTGA-AGGGTCTTGGCGGATAAAGA |
| 4207162 | - | -16 | 1 | parA | sporulation initiation inhibitor protein Soj | AAGATAGTACATGTTCATGT-GAAAGTAGGTGA-CATCGTGGGAAAAATCATAG |
